# The genome of *Drosophila innubila* reveals lineage-specific patterns of selection in immune genes

**DOI:** 10.1101/383877

**Authors:** Tom Hill, Boryana S. Koseva, Robert L. Unckless

## Abstract

Pathogenic microbes can exert extraordinary evolutionary pressure on their hosts. They can spread rapidly and sicken or even kill their host to promote their own proliferation. Because of this strong selective pressure, immune genes are some of the fastest evolving genes across metazoans, as highlighted in mammals and insects. *Drosophila melanogaster* serves as a powerful model for studying host/pathogen evolution. While *Drosophila melanogaster* are frequently exposed to various pathogens, little is known about *D. melanogaster*’s ecology, or if they are representative of other *Drosophila* species in terms of pathogen pressure. Here, we characterize the genome of *Drosophila innubila*, a mushroom-feeding species highly diverged from *D. melanogaster* and investigate the evolution of the immune system. We find substantial differences in the rates of evolution of immune pathways between *D. innubila* and *D. melanogaster*. Contrasting what was previously found for *D. melanogaster*, we find little evidence of rapid evolution of the antiviral RNAi genes and high rates of evolution in the Toll pathway. This suggests that, while immune genes tend to be rapidly evolving in most species, the specific genes that are fastest evolving may depend either on the pathogens faced by the host and/or divergence in the basic architecture of the host’s immune system.

## Introduction

Pathogens are a substantial burden to nearly every species on the planet, providing a strong selective pressure for individuals experiencing infection to evolve resistance. There is considerable evidence for selection acting on genes involved in resistance to this pathogenic burden, as highlighted by several studies across Metazoans (Kimbrell and Beutler 2001; Enard et al. 2016) including *Drosophila* (Sackton et al. 2007; Obbard et al. 2009). Much work concerning the evolution of the invertebrate immune system has focused on the *D. melanogaster.* While pathogen pressure is ubiquitous, the diversity of pathogens that hosts face and the selection for resistance they trigger vary tremendously (Sackton et al. 2007; Hetru and Hoffmann 2009; Sackton et al. 2009; Merkling and van Rij 2013; Martinez et al. 2014; Hanson et al. 2016; Martinson et al. 2017; Palmer, et al. 2018a). So, while it is abundantly clear that immune genes are often among the fastest evolving genes in the genome (Obbard et al. 2006; Sackton et al. 2007; Obbard et al. 2009; Enard et al. 2016; Shultz and Sackton 2019), it is less clear whether the same genes are fast evolving in species that are genetically, geographically and/or ecologically diverged.

Based on genetic work, a number of pathways have been implicated in immune response based on work in *D. melanogaster* (Hoffmann 2003). Three of these pathways are Toll, IMD and JAK-STAT, implicated in the defense response to gram-positive bacteria and Fungi, defense response to Gram-Negative Bacteria, and general stress response respectively (Ekengren and Hultmark 2001; Hoffmann 2003; Hultmark 2003; Hetru and Hoffmann 2009; Darren J. Obbard et al. 2009). In all three pathways, several genes are found to be rapidly evolving, likely due to Host/parasite arms races (Obbard et al. 2006; Sackton et al. 2007; Darren J. Obbard et al. 2009; Sackton et al. 2009). In fact, orthologs of some effector molecules in these pathways are difficult to identify due to their rapid evolution (Ekengren and Hultmark 2001; Sackton et al. 2007; West and Silverman 2018).

In *D. melanogaster* and several members of the *Sophophora* subgenus, the canonical antiviral RNA interference (RNAi) genes are some of the fastest evolving genes in the genome, and are the focus of many studies of the evolution of antiviral pathways (Obbard et al. 2006; Obbard et al. 2009; Palmer et al. 2018a). These studies suggest that viruses, specifically RNA viruses, are a major selective pressure in *D. melanogaster*, requiring a rapid evolutionary response from the host (Obbard et al. 2006; Obbard et al. 2009; Daugherty and Malik 2012; Palmer et al. 2018b). The primary antiviral pathway characterized in *D. melanogaster* is an RNA interference system, which uses small interfering RNAs (siRNA), generated from double stranded viral messenger RNAs (mRNAs) and Argonaute-family proteins (Hutvagner and Simard 2008; Sabin et al. 2009), to bind complimentary sequences and degrade them, preventing their use as a translation template and stopping viral replication (Wang et al. 2006; Obbard et al. 2009; Ding 2010).Importantly, these pathways have mostly been validated only in *D. melanogaster*, therefore, a broader view of antiviral immune gene evolution across *Drosophila* is warranted.

One particular group in the *Drosophila* genus with a rich history of ecological study, and with great potential as a host/pathogen study system, is the quinaria group (Jaenike and Perlman 2002; Perlman et al. 2003; Dyer et al. 2005; Jaenike and Dyer 2008; Unckless 2011; Unckless and Jaenike 2011). This species group is mostly mycophagous, found developing and living on the fruiting bodies of several (sometimes toxic) mushrooms. These mushrooms are commonly inhabited by parasitic nematode worms, trypanosomes and a host of parasitic microbes (Dyer et al. 2005; Martinson et al. 2017) which are likely significant pathogenic burdens, requiring a strong immune response. One member of the quinaria group of particular interest concerning host/pathogen coevolution is *Drosophila innubila* (Patterson and Stone 1949). While many species in the quinaria group are broadly dispersed across temperate forests (including the sister species *D. falleni* and outgroup species *D. phalerata*) (Patterson and Stone 1949; Markow and O’Grady 2006), *Drosophila innubila*, is limited to the “Sky Islands”, montane forests and woodlands southwestern USA and Mexico. It likely colonized the mountains during the previous glaciation period, 10 to 100 thousand years ago (Patterson and Stone 1949; Jaenike et al. 2003). The flies are restricted to elevations of 900 to 1500m, and are active only during the rainy season (late July to September) (Jaenike et al. 2003). *D. innubila* is also the only species in the quinaria group frequently (25-46% in females) infected by a male-killing *Wolbachia* strain (*wInn*), leading to female biased sex-ratios (Dyer 2004). This *Wolbachia* is closely related to *wMel*, which infects *D. melanogaster* (but does not kill males) (Jaenike et al. 2003). Interestingly, *D. innubila* is also frequently (35-56%, n > 84) infected by Drosophila innubila Nudivirus (DiNV), thought to have spread to *D. innubila* during their expansion in the glaciation period (Unckless 2011; Hill and Unckless 2017). In contrast, DiNV is found at lower frequencies in *D. innubila’s* sister species, *D. falleni* (0-3%, n = 95) and undetected in the outgroup species, *D. phalerata* (0%, n = 7) (Unckless 2011). DiNV reduces both lifespan and fecundity of infected hosts (Unckless 2011), with related viruses also causing larval lethality (Payne 1974; Wang and Jehle 2009). When infected with a similar DNA virus (Kallithea virus), *Drosophila melanogaster* show a standard antiviral immune response, including the induction of the antiviral siRNA pathway along with other identified antiviral pathways (Palmer et al. 2018a; Palmer et al. 2018b). Thus, despite lacking the genomic resources of *D. melanogaster*, *D. innubila* and *D. falleni* are potentially useful model systems to understand the evolution of the immune system to identify if the signatures of selection are conserved across pathways between the highly genetically and ecologically diverged melanogaster and quinaria groups. Like *D. melanogaster* (Sackton et al. 2007; Obbard et al. 2009), genes involved in immune defense will be rapidly evolving in this species group. Though the difference in pathogen pressure may affect which pathways are rapidly evolving.

These results suggest species with different pathogenic burdens may have different rates of immune evolution. To this end, we surveyed the evolutionary divergence of genes within a trio of closely related mycophagous *Drosophila* species (*D. innubila, D. falleni* and *D. phalerata*). As a first step, we sequenced and assembled the genome of *D. innubila*, resulting in an assembly which is on par with the *Drosophila* benchmark, *D. melanogaster* release 6 (Dos Santos et al. 2015). Using short read alignments to *D. innubila* of two closely-related species (*D. falleni* and *D. phalerata*), we found evidence of selective constraint on the antiviral RNAi pathway, but rapid evolution of genes in several other immune pathways, including several conserved broad immune pathways such as the Toll and JAK-STAT pathways. These suggest that pathogen pressure differences may lead to drastic differences in immune evolution, or environmental changes may cause differences in general stress response and developmental pathways.

## Results

### Genome sequencing and assembly

*D. innubila* is a mushroom-feeding species found across the sky islands of the South-Western USA and Western Mexico (Patterson and Stone 1949). It is in the quinaria group of the *Drosophila* subgenus, approximately 50 million years diverged from the research workhorse, *D. melanogaster* (Dyer 2004; Markow and O’Grady 2006). *D innubila* has a sister species in northern North America, *D. falleni*, and the pair share an old-world outgroup species, *D. phalerata*. These species are highly diverged from all other genome-sequenced *Drosophila* species and represent a genomically understudied group of the *Drosophila* subgenus (Jaenike et al. 2003; Markow and O’Grady 2006).

We sequenced and assembled the genome of *D. innubila* using a combination of MinION long reads with HiC scaffolding (Oxford Nanopore Technologies Inc., Oxford, UK), and Illumina short reads for error correcting (Illumina, San Diego, CA) (Figure 1A, Table 1; NCBI accession: SKCT00000000). The genome is 168 Mbp with 50% of the genome represented in scaffolds 29.59Mbp or longer (N50=29.59Mbp), eclipsing release 6 of *D. melanogaster* (N50=25.29Mb, (Clark et al. 2007; Dos Santos et al. 2015; Gramates et al. 2017). Additionally, the *D. innubila* N90 is 25.98Mbp compared to 23.51Mbp in *D. melanogaster*. At this quality of assembly, the N50 and N90 statistics are approaching full chromosome lengths in *Drosophila* and therefore likely do not represent how contiguous the assembly is, given the genome of interest. To better discern between the quality of assemblies, we calculated the proportion of the entire genome found in the six largest contigs (for the six *Drosophila* Muller elements) for each reference genome. In *D. innubila*, 97.71% of the genome is found in these six largest contigs, compared to 95.99% of the *D. melanogaster* genome (Supplementary Tables 1-3).

**Figure 1:**
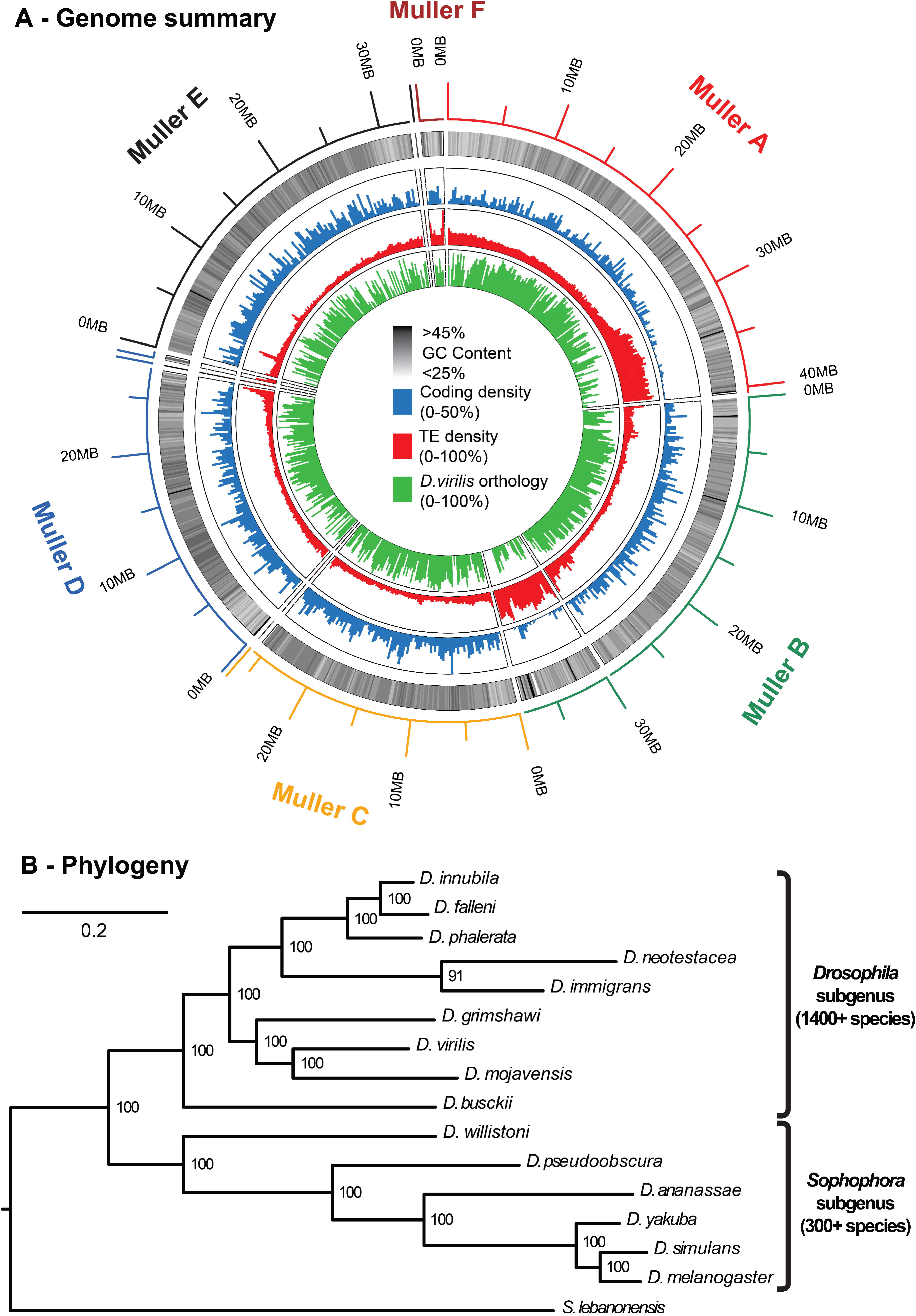
The *D. innubila* genome. **A.** A circular summary of the *D. innubila* genome limited to the assembled acrocentric chromosomes on ten major scaffolds. The rings (from the outside in) are the chromosome identity, the length of each segment, the percentage GC content, the coding density, the transposable element density (250 kilobase pairs windows, sliding 250 kilobase pairs), and the percentage of each window that aligns to *D. virilis* (250kbp windows, sliding 250kbp). **B.** Phylogenetic relationships of *D. innubila*, *D. falleni, D. phalerata* and the main lineages of *Drosophila* generated using 100 concatenated genes, with 500 bootstraps (shown as % support on the nodes).

**Table 1:** Genome Summary. Summary of the major assembled scaffolds in the *D. innubila* genome, including the length in kilobase pairs (kbp), number of genes, number of orphan genes, gene lengths in base pairs (bp) and proportion of the scaffold that is repetitive content.

| Scaffold (Muller element - *D. melanogaster* ortholog - ID) | Total Length (kbp) | Total Gene count | Orphan count | Mean gene length (bp) | Mean repeat content (%) |
| --- | --- | --- | --- | --- | --- |
| A-X-3 | 40479 | 2012 | 73 | 5090.7 | 31.7 |
| B-2L-4 | 29570.5 | 2249 | 99 | 4394.6 | 13.9 |
| B-2L-5 | 8037.1 | 160 | 13 | 4679.2 | 62.9 |
| C-2R-1 | 25683.3 | 2518 | 58 | 4033.3 | 12.2 |
| C-2R-27 | 16.5 | 0 | 0 | NA | 92.4 |
| D-3L-0 | 27707.2 | 2345 | 68 | 4248.1 | 12.6 |
| D-3L-29 | 45.7 | 0 | 0 | NA | 83.7 |
| E-3R-2 | 32746.5 | 2863 | 73 | 4092.4 | 11.4 |
| E-3R-11 | 18.4 | 1 | 1 | 282 | 75.6 |
| F-4-6 | 2027.1 | 106 | 7 | 6992 | 35.5 |
| Unassembled (324 contigs) | 1163.5 | 46 | 1 | 1429 | 39.1 |
| Mitochondria | 16.1 | 18 | 0 | 1658.7 | 0 |
| **Total** | **168031** | **12318** | **393** | **4350.7** | **13.5** |

Using a combined transcriptome assembly of all life stages and protein databases from other species, we annotated the genome and found 12,318 genes (including the mitochondrial genome), with coding sequence making up 11.5% of the genome, at varying gene densities across the Muller elements (Figure 1A blue, Table 1). Of the annotated genes, 11,925 (96.8%) are shared with other *Drosophila* species (among the 12 genomes available on Flybase.org), 7,094 (57.6%) have orthologs in the human genome, and the annotation recovered 97.2% of the Dipteran BUSCO protein library (Simão et al. 2015). The *D. innubila* genome has an average of 36.57% GC content, varying across the Muller elements (Figure 1A black/white). Using dnaPipeTE (Goubert et al. 2015), RepeatModeler (Smit and Hubley 2008) and RepeatMasker (Smit and Hubley 2015), we estimated that 13.53% of the genome consists of transposable elements (TEs, Figure 1A red), which is low for *Drosophila* (Sessegolo et al. 2016). Using Mauve (Darling et al. 2004) we compared our assembly to the best assembled genome within the *Drosophila* subgenus, *D. virilis*, and found that most regions in the *D. innubila* genome have some orthology to the *D. virilis* reference genome (Figure 1A green). A summary of the assembled genome including codon bias, TE content, orphan genes, duplications and expression changes across life stages can be found in the supplementary results (Supplementary Tables 7-20 and Supplementary Figures 4-12).

### Immune pathways evolve differently in D. innubila and D. melanogaster

We aimed to determine whether evolutionary rates in different functional categories are conserved across *D. innubila* and *D. melanogaster*, given the genetic and ecological divergence between the two species (Patterson and Stone 1949; Jaenike et al. 2003; Markow and O’Grady 2006). Using short reads from *D. falleni* and *D. phalerata* mapped to the *D. innubila* genome, we identified DNA sequence divergence and generated consensus gene sequences for each species. We then aligned the DNA sequence from each species for each gene to the *D. innubila* ortholog (PRANK –codon +F) (Löytynoja 2014). For each ortholog set, we identified the proportion of synonymous (*dS*) substitutions and amino acid changing, non-synonymous substitutions (*dN*) (per possible synonymous or non-synonymous substitution respectively) occurring on each branch of the phylogeny (codeML branch based approach, model 0) (Yang 2007; McKenna et al. 2010; DePristo et al. 2011; Löytynoja 2014). This allowed us to calculate *dN/dS* to identify genes showing signatures of rapid or unconstrainted evolution specifically on the *D. innubila* branch of the tree (elevated *dN/dS*, Figure 2A). Conversely, we can also identify genes under strong purifying selection (reduced *dN/dS*) (Yang 2007). We also performed this analysis across the *D. melanogaster* clade using *D. melanogaster*, *D. simulans* and *D. yakuba* reference genomes but focusing on the *D. melanogaster* branch (Clark et al. 2007; Gramates et al. 2017). These trios are of similar levels of divergence (Figure 1B), The rate of evolution, as measured by *dN/dS*, is significantly positively correlated between the species across individual genes (species-specific branch comparison, Spearman rank correlation = 0.6856, *p*-value = 1.52×10^−06^) and most genes are under selective constraint; only 25.6% of genes have *dN/dS* greater than 0.5 (Figure 2B).

**Figure 2:**
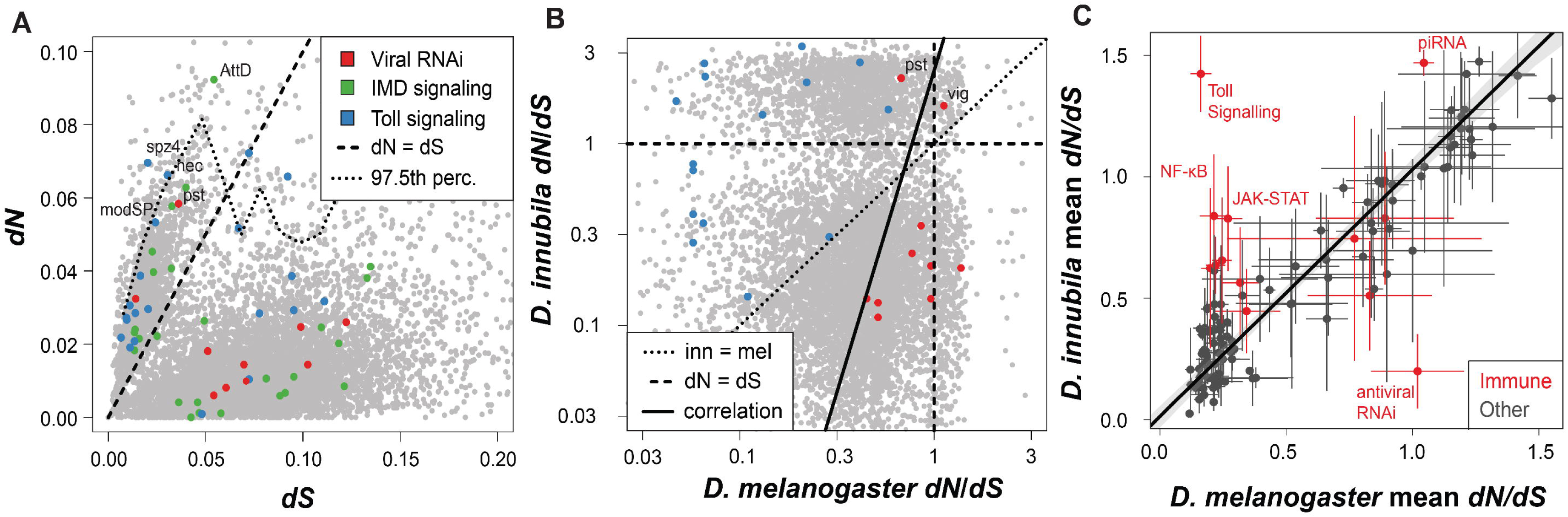
Toll and antiviral genes evolve differently between *D. melanogaster* and *D. innubila*. **A.** The non-synonymous (potentially adaptive) divergence (*dN*) for each gene across the *D. innubila* genome, compared to the gene’s synonymous (neutral) divergence (*dS*). Genes involved in antiviral and antibacterial (*Toll* and *IMD*) immune signaling pathways are colored. The upper 97.5^th^ percentile is shown as a dotted line, *dN/dS* = 1 is shown as a dashed line. **B.** Comparison of *dN/dS* in *D. innubila* and *D. melanogaster*. Toll and antiviral genes are shown for comparison. The dotted line highlights when *innubila* and *melanogaster* dN/dS are equal. The dashed line highlights when dN=dS in either species. The solid line highlights the spearman correlation between *D. melanogaster* and *D. innubila* dN/dS. **C.** Mean *dN/dS* for genes within core 116 GOslim categories (Consortium et al. 2000; Carbon et al. 2017) in both *D. melanogaster* and *D. innubila*, alongside specific immune and RNAi categories of interest. Bars are shown giving the standard error for each category in *D. melanogaster* (X-axis) and *D. innubila* (Y-axis). Categories are colored by immune categories (red) or background categories (grey), with immune categories of interest highlighted. The fitted line is for all background categories, with immune categories added *post hoc*. **Abbreviations:** *AttD* = *attacin D*, *modSP* = *modular serine protease*, *nec = necrotic*, *pst* = *pastrel*, *spz4* = *spaetzle 4*, *vig = vasa intronic gene*.

To determine whether similar classes of genes were under similar selection pressures in the two species, we grouped genes by gene ontology (GO) categories using GOSlim (Consortium et al. 2000; Carbon et al. 2017). There was a significant linear correlation between mean values of GO categories in each species (Figure 2C black points, Spearman rank correlation = 0.847, GLM *t* = 17.01, *p*-value = 4.613×10^−33^), suggesting that, in general, pathways may be under similar selective pressures (either constrained evolution or positive selection, Figure 2C). We identified GO categories enriched in the top 10% of genes for *dN/dS* to find categories evolving rapidly in each species. Several enriched GO immune categories were common to both *D. innubila* and *melanogaster*, including defense response to bacteria and antimicrobial peptide regulation (Supplementary Table 4). However, several GO terms related to Toll signaling were enriched exclusively in *D. innubila*, while gene silencing by RNA and RNA splicing were enriched in *D. melanogaster* but not *D. innubila* (Figure 2B red points, Supplementary Table 4).

We fitted a model to contrast *D. innubila dN/dS* and *D. melanogaster dN/dS* per gene and extracted the studentized residuals. We examined the upper 10% of residuals, which should contain genes fast evolving in only *D. innubila*. We found Toll receptor signaling pathways and metabolic processes enriched, among others (Supplementary Table 4, Figure 2B & C, *p*-value < 0.000775, though this is not significant after correcting more multiple tests). Conversely, in the lower 10% (which should be genes fast-evolving only in *D. melanogaster*) we found RNAi and response to virus genes enriched (Supplementary Table 4, Figure 2B & C, *p*-value < 0.000998, though this is not significant after correcting for multiple tests). These lines of evidence suggested that, while many functional categories are evolving similarly, Toll and antiviral RNAi pathways are evolving quite differently between *D. melanogaster* and *D. innubila* and motivated a more thorough examination of the differences in the evolution of genes involved in immune defense.

### Immune evolution differs between species groups, even after controlling for synonymous divergence

While we found no correlation between *dN/dS* and gene length in either species (GLM *t* = 0.34, *p*-value = 0.81), we did find a significant negative correlation between *dN/dS* and *dS* in *D. innubila* (Figure 2A, GLM *t* = −64.94, *p*-value = 2.2×10^−16^). Most genes with high *dN/dS* had lower values of *dS*, possibly due to the short gene branches (and low neutral divergence) between species inflating the proportion of non-synonymous substitutions (Figure 2A). We also found slightly different distributions of *dN/dS* in each species, suggesting which may cause the differences seen (Supplementary Figure 1). Because of these effects, we attempted to control for differences by extracting genes that were in the upper 97.5% *dN/dS* of genes per 0.01 *dS* window and with *dN/dS* greater than 1 and labeled these 166 genes as the most rapidly evolving on the *D. innubila* branch. Contrasting *D. melanogaster* (Obbard et al. 2006), these genes were not enriched for antiviral RNAi genes and were instead significantly enriched for several metabolic and regulation pathways, as was found previously (Table 2). The most common type of gene with elevated *dN/dS* were those involved in the regulation of the Toll pathway (Table 2, Figure 2 & 3, Supplementary Table 5, GOrilla FDR *p*-value = 0.00015 after multiple testing correction, enrichment = 14.16) (Eden et al. 2009). Specifically, we found four Toll signaling genes; *spatzle4, necrotic, spheroide* and *modSP*; were the fastest evolving genes in this pathway, and among the fastest in the genome (Figure 2A, above the dotted line, Supplementary Table 6). Most of these rapidly evolving genes are signaling genes, which were, as a class, not particularly fast evolving in the *D. melanogaster* clade (Sackton et al. 2007). Additionally, *Pp1apha-96A* and *Attacin D*, genes involved in Gram-negative bacterial response, were rapidly evolving (upper 97.5% of genes, *dN/dS* > 1).

**Figure 3:**
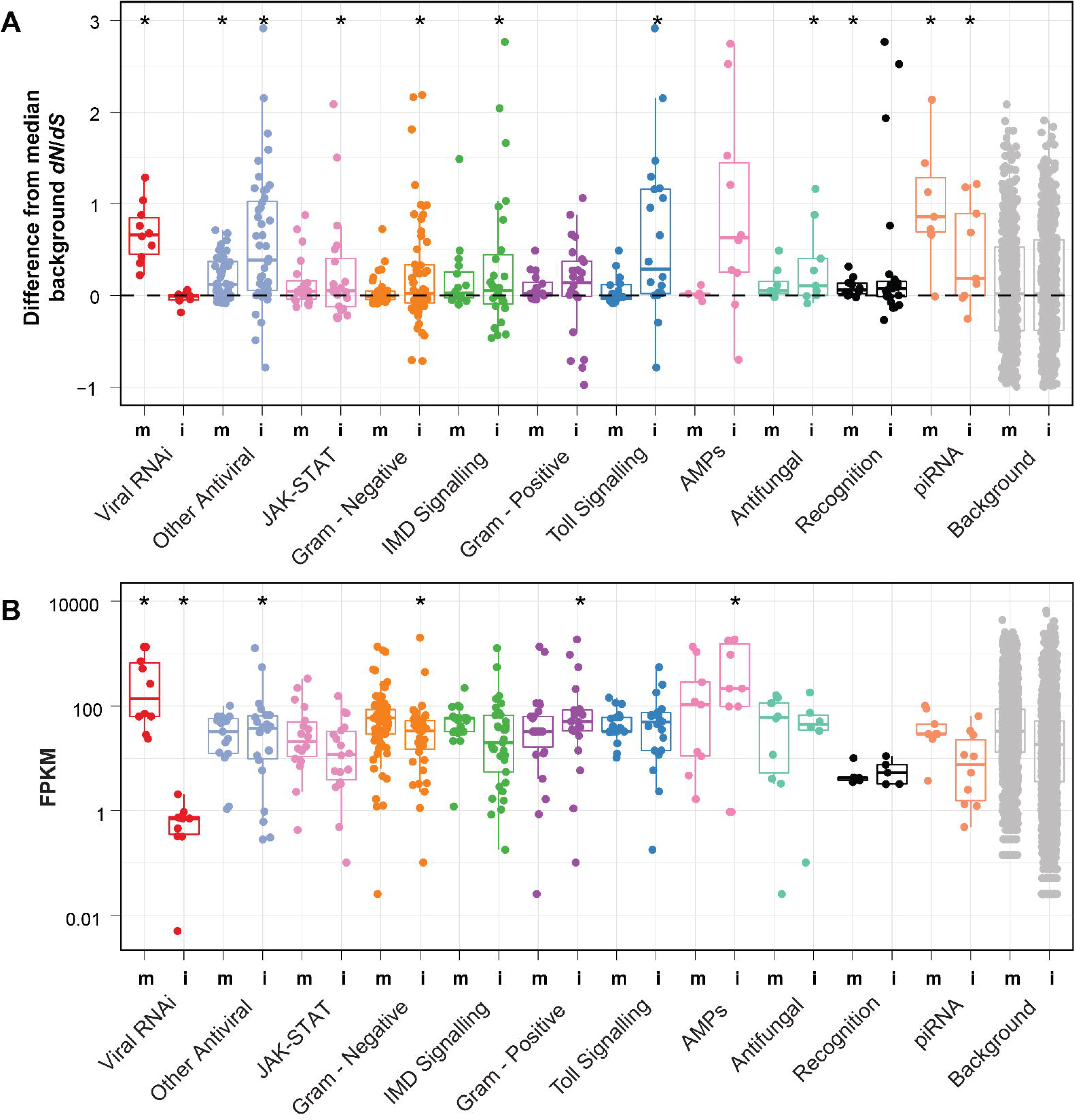
Fast evolving immune gene categories differ between species. **A.** For each immune gene or RNAi gene, we have calculated the difference in *dN/dS* between the gene and the mean of background genes of similar *dS* (+-0.01*dS*). i refers to the *D. innubila* branch while m refers to the *D. melanogaster* branch. A *p*-value (from a single sample, two-sided *t*-test looking for significant differences from 0) of 0.05 or lower is designated with *. **B.** Expression as read counts per 1kbp of exon for each gene (FPKM) by immune gene in each species. For each category we have overlaid a boxplot showing the median (center line) and interquartile range for each category in both species groups, with whiskers to 97.5% of the next interquartile. i refers to *D. innubila* while m refers to *D. melanogaster*. Categories marked with a * are significantly different from the background category with a Mann-Whitney U test (*p*-value < 0.05).

**Table 2:** Gene enrichments. Gene ontology groups enriched for high dN/dS on the *D. innubila* branch. Table includes the number of genes in each pathway found in the upper 0.25% for *dN/dS*, the enrichment of each gene category as well as significance of the category before and after multiple testing correction.

| Gene Ontology category | No. genes<br>dN/dS ><br>0.25% | Total No.<br>genes in<br>GO<br>category | Enrichment | *p*-value | *p*-value (after<br>multiple<br>testing<br>correction) |
| --- | --- | --- | --- | --- | --- |
| Regulation of the Toll<br>Pathway | 4 | 20 | 14.16 | $2.62 \times 10^{-8}$ | 0.000153 |
| Metabolic Process | 160 | 2317 | 1.22 | $6.29 \times 10^{-4}$ | 1 |
| Cellular response to light<br>stimulus | 5 | 25 | 6.77 | $6.98 \times 10^{-4}$ | 1 |
| Vesicle uncoating | 2 | 2 | 33.85 | $8.68 \times 10^{-4}$ | 1 |
| Organic hydroxy<br>compound metabolic<br>process | 9 | 87 | 3.5 | $9.85 \times 10^{-4}$ | 1 |

Given the differences between species, we compared molecular evolution of genes in each of immune gene category for the entire *melanogaster* trio (*D. melanogaster, simulans* and *yakuba*) with the evolution of those same genes in the entire *D. innubila* trio (*D. innubila, falleni* and *phalerata*), alongside comparing specifically *D. innubila* and *D. melanogaster*. Because non-synonymous divergence is elevated in genes with low synonymous divergence on the *D. innubila* branch but not the *D. melanogaster* branch (Figure 2A, Supplementary Figure 1), we attempted to control for its effect. For each focal immune gene, we extracted genes on the same chromosome with *dS* within 0.01 of the focal gene. We then calculated the differences in the median *dN/dS* of these control genes and the focal genes, for each branch on the tree, and categorized these differences by immune category based on Flybase gene ontologies (Gramates et al. 2017). We also separated antiviral genes into those associated with antiviral RNAi and those involved in other pathways (such as NF-κB signaling molecules). Using this method, we found most immune categories had slightly positive differences compared to the controls, suggesting faster evolution than the background (Figure 3A), consistent with results across the entire genus (Sackton et al. 2007). Specifically, the Toll signaling, JAK-STAT, response to Gram-positive infection, response to Gram-negative infection and other genes associated with resistance to viral infection (Magwire et al. 2011; Magwire et al. 2012; Palmer et al. 2018a) were significantly higher than the background in the *D. innubila* trio (Figure 3A, Supplementary Figure 3, Supplementary Table 5, *t*-test *t* = 2.39, *p*-value < 0.05, all categories are normally distributed, Shapiro-Wilk test *p-*value > 0.0521). Toll genes also had significantly higher rates of evolution in *D. innubila* than *D. melanogaster* (Wilcoxon Rank Sum Test, W = 226, *p*-value = 0.01051). Again, in contrast to *D. melanogaster* (Obbard et al. 2006), there was no significant elevation of the rates of evolution in antiviral RNAi genes in *D. innubila*, suggesting selective constraint (*t-*test *t =* 1.0798, *p*-value = 0.3082). In fact, only one antiviral RNAi gene, *pastrel*, appears to be fast-evolving in *D. innubila*, with most genes in this category close to the median *dN/dS* for the *innubila* genome (Figure 2A, 2B & 3A). Interestingly, *pastrel* is among the slowest evolving antiviral gene in *D. melanogaster* (though still in the upper 25% of all genes). Variation in *pastrel* has been associated with survival after *Drosophila* C virus infection (DCV) in *D. melanogaster*, but is not likely involved with antiviral RNAi (Magwire et al. 2011; Magwire et al. 2012; Barbier 2013).

As *dN/dS* may give false signals of rapid evolution due to multiple nucleotide substitutions occurring per site (Venkat et al. 2018), we calculated δ (another measurement of the rate of evolution) using a method that controls for multi-nucleotide substitutions in a single site (Pond et al. 2005; Venkat et al. 2018). δ was broadly positively correlated with *dN/dS* in both species (Spearman correlation = 0.17, *p*-value = 0.0192). Using this method, we corroborated our previous finding that antiviral genes are rapidly evolving exclusively in the *D. melanogaster* trio compared to the background (Supplementary Figure 2, GLM *t* = 4.398, *p*-value = 1.1×10^−05^), while bacterial response genes (both Gram-positive and -negative) are rapidly evolving only in the *D. innubila* trio compared to the background (Supplementary Figure 2, GLM *t* > 2.682, *p*-value < 0.00731).

On some branches of the *D. innubila* trio phylogeny, we find differing signatures from the total *D. innubila* trio phylogeny. Specifically, while Gram-positive bacterial response is fast evolving in *D. falleni*, antifungal and Gram-negative bacterial responses are fast evolving in *D. innubila* (Supplementary Table 5, *t*-test *t* = 2.11, *p*-value < 0.05), whereas none of these three groups are fast evolving in *D. phalerata*. This potentially highlights differences in the pathogens and environments encountered by the three species. Interestingly, Toll signaling, but not Gram-positive defense response, is fast evolving in *D. innubila*, (Figure 3, Supplementary Table 5), suggesting Toll may play a separate role from signaling Gram-positive defense response in *D. innubila*, possibly directed more acutely towards antiviral or antifungal defense (Takeda and Akira 2005; Zambon et al. 2005; Palmer et al. 2018c) or directed towards Toll’s role in the regulation of development (Keshishian et al. 1993; Valanne et al. 2011).

### Several alternative antiviral immune genes are rapidly evolving in both species

We separated the known antiviral pathways and viral interacting genes into specific categories, and examined their evolution in both *D. melanogaster*, *D. innubila* and across each clade to find pathways showing consistent rates of evolution. The JAK-STAT pathway (Janus kinase signal transduction and activation of transcription) is a conserved signaling pathway involved in processes such as immunity, cell death and general stress response, and is implicated in the DNA virus response (Hultmark 2003; West and Silverman 2018). We find this pathway is significantly faster evolving compared to the background across the *D. innubila* trio, while NF-κB, Toll and putative viral-capsid interacting genes are evolving significantly faster than background genes of similar *dS* in both *D. innubila* and *D. melanogaster* trios (Supplementary Figure 3, *t-*test *t* = 2.90, *p*-value < 0.05). Several genes within these categories also showed consistent signatures across the *D. innubila* and *D. melanogaster* trios (Figure 2 & 3, Supplementary Table 4 & 5). Genes rapidly evolving in both lineages include the JAK-STAT cytokines *upd2* and *upd3*, JAK-STAT regulatory genes *CG30423* and *asrij*, the Toll pathway genes *grass* and *GNBP1*, and the NF-κB signaling molecules *relish* and *Aos1*. After controlling for multiple nucleotide substitutions per site with δ (Pond et al. 2005; Venkat et al. 2018), we found that JAK-STAT, NF-κB, Toll and viral capsid associated genes are rapidly evolving in both trios (Supplementary Figure 2, GLM *t* > 3.22, *p*-value > 0.00128), however the putatively viral capsid associated genes are evolving most rapidly in the *innubila* trio (Supplementary Figure 2, GLM *t* = 4.124, *p*-value = 3.73×10^−05^).

### Low antiviral RNAi expression in Drosophila innubila

We examined immunity and RNAi genes in the context of baseline (e.g. constitutive) gene expression in *D. innubila* compared to *D. melanogaster* using ModEncode (Consortium et al. 2011). It should be noted that this is not a well-controlled comparison, whereby differences in expression could be due to different laboratory conditions or other experimental variables as opposed to true baseline expression differences. Nevertheless, some of the observed differences are consistent with rates of molecular evolution. We found no effect for life stage, sex or tissue on immune expression outside of an increase in immune expression when transitioning from embryo to larval stages (GLM p > 0.05, Supplementary Tables 12-19). Specifically, the Toll pathway genes *Toll*, *GNBP1* and *grass* had higher expression in larvae than embryos, and this was maintained throughout the rest of the life stages. Because the main shift in gene expression appears to occur as embryos develop into larvae, we focused on adults, as they represent a more stable period of gene expression. We focused on adult whole-body expression differences between *D. innubila* and *D. melanogaster*. The viral RNAi pathway is mostly shut off in *D. innubila* (Figure 3B, only 2 of 7 genes showed expression greater than 1 read per million counts per kbp of gene in larvae and adults) and had significantly lower expression than the rest of the genome across all life stages (both before and after adjusting for gene length, Wilcoxon rank sum *p*-value < 0.02). In contrast, the piRNA pathway showed appreciable levels of expression and no difference from the background genome at any stage (Figure 3B, Wilcoxon rank sum *p*-value > 0.05). This difference in expression between the antiviral RNAi and piRNA pathways may be due to antiviral genes only being expressed upon infection, though other antiviral genes show no significant difference in expression from the background (Wilcoxon Rank Sum test: W = 27464, *p*-value = 0.1089). Additionally, AMPs, which are highly induced upon infection in insects, here also showed high levels of constitutive expression compared to the background (Figure 3B, Wilcoxon Rank Sum test: W = 11642, *p*-value < 0.05). Furthermore, the antiviral RNAi genes are significantly more highly expressed compared to background genes in the *D. melanogaster* expression data, even without a known infection, and are further induced upon infection with a DNA virus (Wilcoxon Rank Sum test: W = 80672, *p*-value < 0.005) (Obbard et al. 2006; Obbard et al. 2009; Ding 2010; Palmer et al. 2018a; Palmer et al. 2018c). Antiviral RNAi genes are significantly more highly expressed in *D. melanogaster* than in *D. innubila* (Wilcoxon Rank Sum test: W = 2, *p*-value = 0.0003), while immune signaling and immune recognition genes are more highly expressed in *D. innubila* compared to *D. melanogaster* (Wilcoxon Rank Sum test: W = 178 *p*-value = 0.0002). Thus, high rates of expression seem to be associated with high rates of evolution in *Drosophila* immune genes, irrespective of species.

## Discussion

Host/parasite coevolution is ubiquitous across the tree of life (Dawkins and Krebs 1979; Lively 1996). It is expected to result in rapid evolution of protein sequence in both the host and the parasite, as both organisms are adapting to escape the selective pressure exerted by the other. In support of this, immune genes in *Drosophila*, and other organisms, evolve more rapidly than most other gene categories (Kimbrell and Beutler 2001; Sackton et al. 2007; Obbard et al. 2009; Enard et al. 2016), the fastest among these being the antiviral genes (Obbard et al. 2006; Obbard et al. 2009; Enard et al. 2016; Palmer et al. 2018a). Here, we have shown that while immune genes are fast evolving in *D. innubila*, the categories of genes most rapidly evolving is strikingly different from those most rapidly evolving in *D. melanogaster*.

There are several explanations for the observed differences in immune evolution between the *D. melanogaster* and *D. innubila* trios. First, and most obvious explanation is that different pathogen pressures result in different rates of evolution between species. The most tempting difference to highlight is the high frequency DNA virus infection in *D. innubila* but not *D. melanogaster* (Unckless 2011). DNA virus response in *Drosophila* involves a larger set of pathways than RNA virus response, which is largely mediated via the siRNA pathway (Coccia et al. 2004; Bronkhorst et al. 2012; Palmer et al. 2018b). Many previous studies of RNA and DNA virus immune response in *D. melanogaster* have implicated the IMD, JAK-STAT, NF-κB and Toll pathways as vital components of viral defense, all of which are rapidly evolving in *D. innubila* (Dostert et al. 2005; Zambon et al. 2005; Hetru and Hoffmann 2009; William H Palmer et al. 2018; West and Silverman 2018). Due to overlapping viral transcripts, infection can still induce the siRNA pathway irrespective of the viral class, but may differ in effectiveness between species and viral class (Webster et al. 2015; Palmer et al. 2018c). It is currently believed that *D. melanogaster* is mostly exposed to RNA viruses in nature whereas *D. innubila* is mostly exposed to DNA viruses (Unckless 2011; Webster et al. 2015; Lewis et al. 2018). Therefore, it makes sense that various immune response pathways are evolving at different rates in the two species groups. In keeping with this, *D. falleni* and *D. phalerata* have different rates of immune gene evolution and are not frequently exposed to the DNA virus (Unckless 2011) (Supplementary Tables 5 and 6).

DNA virus exposure is not the only difference in pathogens seen between *D. innubila* and *D. melanogaster*. In fact, most pathways identified as rapidly evolving in *D. innubila* are involved in the response to infection by multiple pathogens (Toll, for viruses, Fungi and Gram-positive bacteria) or are general stress response pathways (JAK-STAT). Given the similar ecologies (including regular exposure to rotting mushrooms) (Jaenike et al. 2003; Perlman et al. 2003) and consistent signatures of rapid evolution in *D. innubila* and *D. falleni*, similar pathogen pressures may drive rapid evolution of these pathways in quinaria group flies (Shoemaker et al. 1999; Perlman et al. 2003). For example, the rapid evolution of Toll signaling but not Gram-positive defense response (Figure 3), might suggest that Toll is evolving in response to something other than Gram-positive bacteria such as fungal pathogens, viruses or even extracellular parasites that uniquely infect the quinaria group (Jaenike and Perlman 2002; Hoffmann 2003; Perlman et al. 2003; Zambon et al. 2005; Hamilton et al. 2014; Palmer et al. 2018b).

Previous work in *D. melanogaster* has highlighted the role that Gram-negative commensal bacteria play in priming the antiviral immune system (Sansone et al. 2015). As both the Toll and IMD signaling pathways are rapidly evolving across the *innubila* trio, it is conceivable that this priming may play a role in immune defense in *D. innubila*.

Finally, Toll signaling is also involved in dorso/ventral development and motorneuron development in *Drosophila* (Keshishian et al. 1993; Hoffmann 2003; Valanne et al. 2011), as eye and neuronal development are almost always enriched in *D. innubila* but not *D. melanogaster* (Supplementary Table 4), this rapid evolution of Toll may have little to do with immune response and instead is involved in changes in the body pattern to adapt to changes in the environment of *D. innubila*. Though this explanation does not explain the rapid evolution of other immune pathways.

A second hypothesis for the lack of evolution in the antiviral RNAi system, is that the immune response to DNA virus infection has diverged in the approximately 50 million years since the quinaria and melanogaster groups last shared a common ancestor. They may fundamentally differ in their immune response to viral infection, and this may be due to the divergence of the siRNA pathways across the *Drosophila* groups (Lewis et al. 2018). This divergence could also be non-adaptive, in fact, given *D. innubila* has undergone repeated bottlenecking during habitat invasion, it is possible that changes in effective population size may have led to genetic drift steering the evolution of the immune system in this species, resulting in relaxed constraint on immunity genes. An ineffective antiviral immune system may even explain the high frequency of DiNV infection in *D. innubila* (Unckless 2011). However, rates of evolution are mostly consistent across the *D. innubila* trio, and, as broadly dispersed temperate species, *D. phalerata* and *falleni* should not have been affected by the same demographic patterns (Markow and O’Grady 2006). *D. innubila*’s invasion of the ‘sky islands’ is also estimated to be more ancient than *D. melanogaster*’s global invasion (occurring during the last glaciation period, 10-100KYA), with current estimates of diversity at similar levels as *D. melanogaster* (Dyer and Jaenike 2005). If *D. innubila’s* bottleneck was more severe than *D. melanogaster*’s, drift may still explain the lack of antiviral evolution in *D. innubila*. The lack of adaptation of the antiviral RNAi could also be due to the more recent infection by DiNV, estimated to have infected *D. innubila* 10-30 thousand years ago, however this is more than enough time for adaptation to occur in *Drosophila* (Aminetzach et al. 2005; Karasov et al. 2010).

There are several other aspects of the host biology that may explain the constrained evolution of the siRNA in *D. innubila* (Figure 2, 3). siRNA, alongside piRNA have been implicated in transposon regulation as well as viral suppression (Biryukova and Ye 2015). It is possible siRNA has an alternate, TE related role in *D. innubila*, which may contribute to their low TE content (Figure 1, Supplementary Figure 6).

Studies have also identified an interaction between *Wolbachia* infection and resistance or susceptibility to viral infection (Teixeira et al. 2008; Martinez et al. 2014). The high frequency of *Wolbachia* infection in *D. innubila* (Dyer and Jaenike 2005), may therefore provide some resistance to viral infection or may be involved in immune system priming (Sansone et al. 2015). However, *Wolbachia* has only been shown to protect against RNA viruses (Teixeira et al. 2008), and this effect of *Wolbachia* was found to be absent in an earlier assessment of DiNV infections in *D. innubila* (Unckless 2011; Martinez et al. 2014), suggesting *Wolbachia* may not play a role in viral resistance.

We have worked to bring mycophagous *Drosophila* to the table as a modern genomic model for the study of immune system evolution. Here we have highlighted that the evolution of the immune system among closely related trios of species may differ drastically across *Drosophila* genera. Specifically, we found that across the *D. innubila* genome, even though the immune system is, in general, evolving rapidly; the canonical antiviral RNAi pathways do not appear to be evolving as if in an arms race with viruses. Instead, several alternative immune pathways may be evolving in response to the different pathogen pressures seen in this species. Together these results suggest that the evolution of genes involved in the immune system can be quite specific to the suite of pathogens faced by hosts.

## Methods

### DNA/RNA isolation, library preparation, genome sequencing and assembly

We extracted DNA following the protocol described in (Chakraborty et al. 2017) for *D. innubila, D. falleni* and *D. phalerata* as further described in the supplementary materials. We prepared the *D. innubila* DNA as a sequencing library using the Oxford Nanopore Technologies Rapid 48-hour (SQK-RAD002) protocol, which was then sequenced using a MinION (Oxford Nanopore Technologies, Oxford, UK; NCBI SRA: SAMN11037163) (Jain et al. 2016). The same DNA was also used to construct a fragment library with insert sizes of ∼180bp, ∼3000bp and ∼7000bp, we sequenced this library on a MiSeq (300bp paired-end, Illumina, San Diego, CA, USA; NCBI SRA: SAMN11037164). We prepared the *D. falleni* and *D. phalerata* samples as Illumina libraries like *D. innubila* but with a 300bp insert size. We sequenced the *D. falleni* fragment library on one half of a MiSeq (300bp paired-end, Illumina, San Diego, CA, USA; NCBI SRA: SAMN11037165) by the KU CMADP Genome Sequencing Core. We sequenced the *D. phalerata* fragment library on a fraction of an Illumina HiSeq 4000 run (150bp paired end, Illumina, San Diego, CA; NCBI SRA: SAMN11037166).

For gene expression analyses, we obtained two replicate samples of female and male heads and whole bodies (including heads), embryos, larvae (pooled across all three instar stages) and pupae (all non-adults were unsexed). RNA was extracted using a standard Trizol procedure (Simms et al. 1993) with a DNAse step. RNA-sequencing libraries were constructed using the standard TruSeq protocol (McCoy et al. 2014) with ½ volume reactions to conserve reagents. Individually indexed RNA libraries (2 replicates from each tissue/sex) were sequenced on one lane of an Illumina “Rapid” run with 100bp single-end reads (NCBI SRA: SAMN11037167-78). All data used in the assembly and annotation of the *D. innubila* genome are available in the NCBI BioProject PRJNA524688.

Bases were called *post hoc* using the built in read_fast5_basecaller.exe program with options: –f FLO-MIN106 –k SQK-RAD002 –r–t 4. Raw reads were assembled using CANU version 1.6 (Koren et al. 2016) with an estimated genome size of 150 million bases and the “nanopore-raw” flag. We then used Pilon to polish the genome with our Illumina fragment library (Walker et al. 2014). The resulting assembly was submitted to PhaseGenomics (Seattle, WA, USA) for scaffolding using Hi-C and further polished with Pilon for seven iterations. With each iteration, we evaluated the quality of the genome and the extent of improvement in quality, by calculating the N50 and using BUSCO to identify the presence of conserved genes in the genome, from a database of 2799 single copy Dipteran genes (Simão et al. 2015).

Repetitive regions were identified *de novo* using RepeatModeler (engine = NCBI) (Smit and Hubley 2008) and RepeatMasker (-gff –gcalc –s) (Smit and Hubley 2015).

These sequencing and assembly steps are further described in the supplementary methods, alongside additional steps taken to verify genes, identify additional contigs and genes, and find genes retained across all species. The final version of the genome and annotation is available on the NCBI (accession: SKCT00000000).

### Genome annotation

As further described in the supplementary methods, we assembled the transcriptome, using all Illumina RNA reads following quality filtering, using Trinity (version 2.4.0) (Haas et al. 2013), Oases (velvetg parameters: -exp_cov 100 -max_coverage 500 -min_contig_lgth 50 -read_trkg yes) (Schulz et al. 2012), and SOAP*denovo* Trans (127mer with parameters: SOAPdenovo-Trans-127mer -p 28 -e 4 and the following kmers: 95, 85, 75, 65, 55, 45, 35, 29, 25, 21) (Xie et al. 2014) which we combined using EvidentialGene (Gilbert 2013; http://eugenes.org/EvidentialGene/). We used the *D. innubila* transcriptome as well as protein databases from *M. domestica*, *D. melanogaster*, and *D. virilis*, in MAKER2 (Holt and Yandell 2011) to annotate the *D. innubila* genome, including using *RepeatModeler* (Smit and Hubley 2008) to not mis-assign repetitive regions. This was repeated for three iterations to generate a GFF file containing gene evidence generated by MAKER2 (Holt and Yandell 2011) (NCBI:).

### Drosophila quinaria group species on the Drosophila phylogeny

To build a phylogeny for the *Drosophila* species including *D. innubila, D. falleni* and *D. phalerata*, we identified genes conserved across all *Drosophila* and humans and found in the Dipteran BUSCO gene set (Simão et al. 2015). We then randomly sampled 100 of these genes, extracted their coding sequence from our three focal species and 9 of the 12 *Drosophila* genomes (Limited to nine due to our focus on the *Drosophila* subgenus and the close relation of several species, rendering them redundant in this instance, Clark *et al*. 2007). We also searched for genomes in the *Drosophila* subgenus with easily identifiable copies of these 100 conserved genes, settling on *D. busckii* (Zhou and Bachtrog 2015)*, D. neotestactea* (Hamilton et al. 2014)*, D. immigrans* and *Scaptodrosophila lebanonensis* (Zhou et al. 2012). We aligned these genes using MAFFT (--auto) (Katoh et al. 2002), concatenated all alignments and generated a phylogeny using PhyML with 500 bootstraps (-M GTR, -Gamma 8, -B 500) (Guindon et al. 2010).

### Signatures of adaptive molecular evolution among species

We mapped short read sequencing data of *D. innubila, D. falleni* and *D. phalerata* to the repeat-masked *D. innubila* reference genome using BWA MEM (Li and Durbin 2009). As similar proportions of reads mapped to the genome (97.6% for *D. innubila*, 96.1% for *D. falleni* and 94.3% for *D. phalerata*), covering a similar proportion of the reference genome (99.1% for *D. innubila*, 98.5% for *D. falleni* and 97.1% for *D. phalerata*), we considered the *D. innubila* genome to be of similar enough to these species to reliably call single nucleotide polymorphisms and indels. We realigned around indels using GATK IndelRealigner then called variants using HaplotypeCaller (default parameters) (McKenna et al. 2010; DePristo et al. 2011). We then used GATK FastaReferenceMaker (default parameters) to generate an alternate, reference genome for each of these species (McKenna et al. 2010; DePristo et al. 2011). We extracted the coding sequence for each gene found in the genomes of *D. innubila, D. phalerata* and *D. falleni* and aligned orthologs using PRANK (-codon +F –f=paml) (Löytynoja 2014). For each PRANK generated gene alignment and gene tree, we used codeML (Yang 2007) for the branches model (M0 model), to identify genes with signatures of rapid evolution on the *D. innubila, D. falleni, D. phalerata* branches, as well as across the entire clade. Focusing specifically in the *D. innubila* branch, for genes involved in the immune system pathways, we attempted to rescale for synonymous divergence. For each focal gene, we found genes with *dS* within 0.01 of the focal gene on the same scaffold. We then found the difference in *dN/dS* between the focal gene and the median of the control gene group.

For an independent contrast, we downloaded the latest coding sequences for *D. melanogaster, D. simulans* and *D. yakuba* from Flybase.org (Downloaded January 2018, Gramates et al. 2017) and aligned orthologous genes using PRANK (-codon +F –f=paml) (Löytynoja 2014). Following the generation of a gene alignment and gene tree, we used codeML (Yang 2007) to identify genes with adaptive molecular signatures on each branch of the phylogeny (using the branch based model, M0). Again, we found the difference in *dN/dS* from background genes of similar *dS* (with 0.01) on the same scaffold for all immune genes, focusing on the *D. melanogaster* branch. We compared genes enriched in the top 2.5% for dN/dS (versus the lower 97.5%) using GOrilla (Eden et al. 2009) in both *D. innubila* and *D. melanogaster*. We also performed this analysis using the top 5% and 10% and found no differences in enrichments than the more stringent 2.5% (not shown).

We downloaded genes involved in a core set of gene ontologies from GOslim (Consortium et al. 2000; Carbon et al. 2017) and found the mean and standard error of *dN/dS* for each category in both *D. melanogaster* and *D. innubila*. We chose to compare genes in the top 10% for dN/dS in both species in these categories, as no enrichments are found in the top 2.5% or 5% for the GOslim genes alone, instead we chose to broadly examine the fastest evolving genes in each species, even if not significantly enriched.

Finally, to control for possibly multiple nucleotide substitutions in a single site creating false signals of rapid evolution (Venkat et al. 2018), we calculated δ using HyPhy (Pond et al. 2005) based on the method presented in (Venkat et al. 2018). δ was calculated under both the null and alternative models, with the best model selected based on the result of a χ^2^ test. We then compared δ between each species and across immune gene categories.

### RNA differential expression analysis

We used GSNAP (-sam –N 1) (Wu and Nacu 2010) to map each set of *D. innubila* RNA sequencing short reads to the repeat masked *D. innubila* genome with the TE sequences concatenated at the end (NCBI SRA: SAMN11037167-78). We then counted the number of reads mapped to each gene per kb of exon using HTSeq (Anders et al. 2015) for all mapped RNA sequencing data and normalized by counts per million per dataset. Mapped RNA sequencing information for *D. melanogaster* across all life stages was downloaded from ModEncode (modencode.org) (Chen et al. 2014). We compared *D. melanogaster* data to *D. innubila* data using EdgeR (Robinson et al. 2009) to identify differentially expressed genes, and also compared RNAseq reads per million reads per 1kbp of exon (fragments per kilobase of exon per million reads, FPKM) between the immune genes of *D. innubila* and *D. melanogaster*.

### Statistical analysis

All statistical analyses were performed using R (R Core Team 2013). We used the R packages EdgeR (Robinson et al. 2009), RCircos (Zhang et al. 2013) and ggplot2 (Wickham 2009) for statistical analysis and plot generation/visualization.

## Supporting information

Supplementary Figures and results

## Acknowledgements

We are thankful to Justin Blumenstiel, Jamie Walters, John Kelly, Carolyn Wessinger, Joanne Chapman and three anonymous reviewers for helpful discussion and advice concerning the manuscript. We thank Brittny Smith and the KU CMADP Genome Sequencing Core (NIH Grant P20 GM103638) for assistance in genome isolation, library preparation and sequencing. Genome annotation was performed in the K-INBRE Bioinformatics Core supported by P20 GM103418. This work was supported by a postdoctoral fellowship from the Max Kade foundation (Austria) and a K-INBRE postdoctoral grant to TH (NIH Grant P20 GM103418). This work was also funded by NIH Grants R00 GM114714 and R01 AI139154 to RLU.

## Supplementary Tables and Figures

**Supplementary Table 1:** Summary of reads used for genome sequencing, assembly, annotation and *dN/dS* calculation.

**Supplementary Table 2:** Summary statistics for each iteration of the genome.

**Supplementary Table 3:** Summary of the genic characteristics of the *D. innubila* genome.

**Supplementary Table 4:** Genes ontologies (GO) enriched for genes with high/low residuals for *dN/dS* between *D. melanogaster* and *D. innubila*, due to drastic differences between the species. Enriched categories are categories which are slow evolving in one species, but fast evolving in the other.

**Supplementary Table 5:** Summary of *dN/dS* statistics for each immune gene category across the total group and on each branch. Additionally, the t-score and p-value for a two-sided *t*-test (μ = 0) for that category is shown. Significant categories are highlighted in bold.

**Supplementary Table 6:** *dN/dS* enrichment for *Drosophila innubila* trio for processes, components and functions, including any enrichments for specific branches.

**Supplementary Table 7:** GO enrichment for processes, components and functions for differential expression between *D. innubila* males and females.

**Supplementary Table 8:** GO enrichment for processes, components and functions for differential expression between *D. innubila* embryos and larvae.

**Supplementary Table 9:** GO enrichment for processes, components and functions for differential expression between *D. innubila* larvae and pupae.

**Supplementary Table 10:** GO enrichment for processes, components and functions for differential expression between *D. innubila* pupae and adults.

**Supplementary Table 11:** *dN/dS* GO enrichment for duplications for processes, components and functions, including any enrichments for specific branches.

**Supplementary Table 12:** A table summarizing the differential gene expression shown in Supplementary Tables 13-19, showing the number of genes differentially expressed between *D. innubila* and *D. melanogaster* at differing life stages, with enrichments in gene ontology (GO) categories.

**Supplementary Table 13:** GO enrichment for processes, components and functions for differential expression between *D. melanogaster* and *D. innubila* embryos.

**Supplementary Table 14:** GO enrichment for processes, components and functions for differential expression between *D. melanogaster* and *D. innubila* larvae.

**Supplementary Table 15:** GO enrichment for processes, components and functions for differential expression between *D. melanogaster* and *D. innubila* pupae.

**Supplementary Table 16:** GO enrichment for processes, components and functions for differential expression between *D. melanogaster* and *D. innubila* adults.

**Supplementary Table 17:** GO enrichment for processes, components and functions for differential expression between *D. melanogaster* and *D. innubila* adult males.

**Supplementary Table 18:** GO enrichment for processes, components and functions for differential expression between *D. melanogaster* and *D. innubila* adult females.

**Supplementary Table 19:** GO enrichment for processes, components and functions for differential expression between *D. melanogaster* and *D. innubila* total samples.

**Supplementary Table 20:** Enrichment or depletion of genes differentially expressed between male and female samples on each scaffold/Muller element.

**Supplementary Figure 1:** Histograms of dN/dS for *D. innubila* and *D. melanogaster*.

**Supplementary Figure 2:** δ (calculated using HyPhy) by immunity category for both *D. innubila* and *D. melanogaster*.

**Supplementary Figure 3:** Difference between viral RNAi, JAK-STAT (filled dots = regulatory, empty dots = cytokines), NF-κB, Toll and putatively viral-interacting proteins from the background *dN/dS* of genes of similar *dS* (+-0.01*dS*) for the *D. melanogaster* branch, the *D. innubila* branch, the total *D. melanogaster* tree and the total *D. innubila* tree. Genes known to be associated with the immune response to viral infection, but no known pathway are classed as ‘Other Antiviral’. A *p*-value (from a two-sided *t*-test looking for significant differences from 0) of 0.05 or lower is designated with *.

**Supplementary Figure 4:** Codon bias distributions across the *Drosophila innubil*a genome, separated by scaffold. CAI = Codon adaptation index. CBI = Codon bias index. Fop = Frequency of optimal codons. GC = Proportion of GC across each gene.

**Supplementary Figure 5:** Comparison between orphan genes and previously described genes, including: **A.** Codon adaptation index (CAI). **B.** Codon bias index (CBI). **C.** Frequency of optimal codons (Fop). **D.** Gene length (in bp). **E.** Number of introns per gene. **F.** Mean expression across life stages (read counts per million).

**Supplementary Figure 6: A.** The proportion of the *D. innubila* genome masked by each type of repeat. LINE = Long interspersed nuclear element RNA transposon, LTR = long terminal repeat RNA transposon, RC = rolling circle DNA transposon, TIR = terminal inverted repeat DNA transposon. **B.** TE content of *D. innubila*, *falleni* and *phalerata*, **C.** Copy number comparisons between *D. innubila*, *D. falleni* and *phalerata*. **D.** dnaPipeTE estimates of the genomic proportion of repetitive elements for each species examined here. Other, NA and SINE categories were removed due to small proportions. Though unlabeled, rRNA is shown in yellow and constitutes 1-2% of the genome.

**Supplementary Figure 7:** Number of TE families found in *D. innubila*, closely related to known TE families (taken from Repbase) in different species group, identified using BLAST, suggesting relatively recent horizontal transfer events.

**Supplementary Figure 8:** *dN/dS* versus *dS* across paralogs for recently duplicated genes. Metal ion binding, protein metabolism and immunity genes are highlighted.

**Supplementary Figure 9:** Volcano plots showing differential gene expression between *D. innubila* and *D. melanogaster* at different life stages. Dots are colored by their significance and if a recent duplication or not (duplicates layered on top), the significance cut off is set a 0.05 following multiple testing correction.

**Supplementary Figure 10:** Volcano plot showing differential gene expression between *D. innubila* male and female samples and significant differences, highlighting if genes are duplicated relative to *D. virilis* or not, and if genes are involved in sperm motility.

**Supplementary Figure 11:** Inversions identified between *D. innubila and D. falleni, and between D. innubila/falleni and D. phalerata* using Pindel (Ye et al. 2009) and Manta (Chen et al. 2016) (taking the consensus of the two programs). Scaffolds are labelled and colored by the Muller element they belong to.

**Supplementary Figure 12:** Size and number of each structural variant between *D. innubila* and *D. falleni* identified using Pindel and Manta (taking the consensus of the two programs).

