## Supplementary Figures and results for "The genome of *Drosophila innubila* reveals lineage-specific patterns of selection in immune genes"

**Alternative antiviral immune pathways are rapidly evolving in *Drosophila innubila***

***Supplementary Methods & Results***

*DNA/RNA isolation, library preparation and sequencing*

We extracted DNA following the protocol described in Chakraborty and Emerson (Chakraborty *et al.* 2017). Briefly, approximately 320 adult females from an isofemale line of *D. innubila* (captured in the Chiricahua Mountains in 2005 by Kelly Dyer, strain name SWRS2005-50) were starved for five hours then frozen and ground to a powder in liquid nitrogen then extracted using a modified version of the Qiagen Blood and Cell Culture DNA Midi Kit (#13343, USA Qiagen Inc., Germantown, MD, USA). The extraction yielded fragment sizes greater than 60,000 bp as determined by Agilent TapeStation (Agilent, Santa Clara, CA, USA). We prepared a sequencing library using the Oxford Nanopore Technologies Rapid 48-hour (SQK-RAD002) protocol which was sequencing using a MinION (Supplementary Table 1, NCBI SRA: SAMN11037163, Oxford Nanopore Technologies, Oxford, UK). The same DNA was also used to construct a Nextera fragment library with insert sizes of ~180bp, ~3000bp and ~7000bp. We sequenced the libraries on a MiSeq (300bp paired-end, Illumina, San Diego, CA, USA) which generated 20104299 200bp paired-end reads (NCBI SRA: SAMN11037164). All data used in the assembly and annotation of the *D. innubila* genome are available in the NCBI BioProject PRJNA524688.

For *D. innubila* long reads, DNA was sequenced on the Oxford Nanopore Technologies Minion platform using the SQK-RAD002 protocol and a 48-hour run (Jain *et al.* 2016). Bases were called *post hoc* using the built in read_fast5_basecaller.exe program with options: –f FLO-MIN106 –k SQK-RAD002 –r–t 4. The MinION produced 746229 reads, an average of 5754bp long, with 656860 reads greater than 1kbp, 225704 reads greater than 10kbp and a maximum read length of 1.61Mbp (NCBI SRA: SAMN11037163).

DNA for the Hi-C protocol was extracted from fifteen adult females by PhaseGenomics with a 4-cutter Sau3AI being used to digest the chromatin. This library was then sequenced on an Illumina NextSeq (Illumina, San Diego, CA, USA).

For the *Drosophila falleni* (strain 15130-1961.00 from the Cornell *Drosophila* species stock center), we followed the same protocol as *D. innubila* for DNA isolation and library preparation, but only constructed a single 300bp insert library. This was sequenced on one half of a MiSeq (300bp paired-end, Illumina, San Diego, CA, USA) by the KU CMADP genomics core. For

*Drosophila phalerata* (obtained from Kelly Dyer), we followed a standard Puregene Gentra extraction (USA Qiagen Inc., Germantown, MD, USA) and constructed a 300bp insert Nextera library (see above) which was sequenced on a fraction of an Illumina HiSeq 4000 run (150bp paired end). This generated 8080281 300bp paired-end reads for *D. falleni* and 24896114 150bp paired-end reads for *D. phalerata*. We estimated the heterozygosity of each sample using Jellyfish (Marcais 2011) and GenomeScope (Vurture *et al.* 2017) and found the heterozygosity of each sample to be between 0.46% and 0.81%.

For gene expression analyses, we obtained two replicate samples of female and male heads and whole bodies (including heads), embryos, larvae (pooled across all three instar stages) and pupae (all non-adults were unsexed). RNA was extracted using a standard Trizol procedure (Simms *et al.* 1993) with a DNAse step. RNA-sequencing libraries were constructed using the standard TruSeq protocol (McCoy *et al.* 2014) with ½ volume reactions to conserve reagents. Individually indexed RNA libraries (2 replicates from each tissue/sex) were sequenced on one lane of an Illumina "Rapid" run with 100bp single-end reads, as outlined in Supplementary Table 1.

*Whole genome assembly*

Raw reads from the Oxford Nanopore Minion were assembled using CANU version 1.6 (Koren *et al.* 2016) with an estimated genome size of 150 million bases and the "nanopore-raw" flag. We then used Pilon (Walker *et al.* 2014) to polish the genome with our Illumina fragment library (default parameters). The resulting assembly was submitted to PhaseGenomics (phasegenomics.com, Seattle, WA USA) for scaffolding using Hi-C and further polished with Pilon for seven iterations. With each iteration, we evaluated the quality of the genome and the extent of improvement in quality, by calculating the N50 and using BUSCO (Simão *et al.* 2015) to identify the presence of conserved genes in the genome, from a database of 2799 single copy Dipteran genes. The final genome and annotation are available at NCBI (accession: SKCT00000000).

*Genome annotation*

We assembled a *de novo* transcriptome using Trinity (version 2.4.0) (Haas *et al.* 2013). First, we quality filtered single-end reads (samples described above) with Scythe (Buffalo 2018; http://github.com/vsbuffalo/scythe) and Sickle (Joshi and Fass 2011;

http://github.com/najoshi/sickle) to remove rRNA, Illumina adapters and low quality sequences (quality less than 20). We concatenated all reads from all tissues and used the Trinity package with default parameters to assemble the transcriptome. We also assembled transcriptome with Oases (Schulz *et al.* 2012) (velvetg parameters: -exp_cov 100 -max_coverage 500 -min_contig_lgth 50 -read_trkg yes) and SOAP*de novo* Trans (Xie *et al.* 2014) (127mer with parameters: SOAPdenovo-Trans-127mer -p 28 -e 4 and the following kmers: 95, 85, 75, 65, 55, 45, 35, 29, 25, 21). These assemblies were used to make a metatranscriptome using EvidentialGene (Gilbert 2013; http://eugenes.org/EvidentialGene/) (parameters: -NCPU=28 -MAXMEM=489000).

Using the *D. innubila* transcriptome as well as protein databases from *M. domestica*, *D. melanogaster*, and *D. virilis*, we searched for evidence of genic regions in the genome assembly. A database containing repeat sequences discovered by RepeatModeler (Smit and Hubley 2008) was also utilized by MAKER2 (Holt and Yandell 2011) to ensure that repetitive regions are not annotated as genes. Post completion of the first MAKER2 run, we extracted all gene models from the annotation that had a predicted protein length of at least 50 amino acids [-l 50] and an AED score (Eilbeck *et al.* 2009) of no more than 0.25 [-x 0.25] to form a training set for SNAP (Korf 2004). The resulting HMM file was used as an input to round 2 of the annotation pipeline, along with GFF files containing all transcript, protein, and repeat evidence collected during round 1. Additionally, we provided MAKER2 with training files for *D. melanogaster* for Augustus (publicly available and distributed with MAKER2) (Stanke *et al.* 2008). After the second round of annotations, we repeated the SNAP training steps taken after round 1, which produced a new HMM file. The HMM file from round 2, the GFF files with the evidence from round 1 (transcripts/proteins/repeats), and the D. melanogaster training set for Augustus were the inputs for round 3 of the annotation pipeline.

Using resources on FlyBase (Gramates *et al.* 2017; FlyBase.org) we identified conservation of each gene by counting the number of the 12 *Drosophila* species genome orthologs (and humans, if applicable). We also calculated the percentage of genic nucleotides per Megabase across the genome in 250kbp sliding windows.

Our annotation resulted in the identification of 12318 genes of varying lengths (Supplementary Table 2 & 3). We find an absence of many tRNAs usually found in *Drosophila* genomes, this may be an error of genome assembly or annotation, but additional tRNAs were unable to be found via BLAST (Altschul *et al.* 1990), either due to their absence in the genome or divergence of tRNAs due to *D. innubila*'s extensive divergence from previously sequenced species (Supplementary Table 3, Figure 1). Most of the genes found in the genome (11925) are shared with other species (among the 12 genomes available on Flybase as of July 2018; ftp://ftp.flybase.net/releases/current), with these genes containing 97.9% of the Dipteran BUSCO library (Simão *et al.* 2015), and 7,094 of these genes have orthologs in the human genome (based on the current version available in FlyBase as of July 2018 ; ftp://ftp.flybase.net/releases/current).

*Further genome assembly*

To identify additional genes missed in the Hi-C assembly, we also took all unmapped reads and assembled these using SPAdes (Bankevich *et al.* 2012). We mapped MiSeq information to the 15587 SPAdes assembled contigs and kept contigs with similar coverage to the CANU assembled scaffolds (25-35 fold coverage) and with Blastn (Camacho *et al.* 2009) hits to known Dipteran sequences (e-value < 0.001), retaining an additional 302 contigs (336 total).

Finally, we used Mauve (Darling *et al.* 2004) to identify regions of orthology between the *Drosophila virilis* genome (Clark *et al.* 2007) and the *D. innubila* genome. We calculated the GC content and percent of windows with identifiable orthology to *virilis,* in 250kbp windows across the *D. innubila* genome using bedTools (Quinlan and Hall 2010).

To assemble the mitochondrial genome, we took a subset (100000 read pairs) of the short-read data generated by MiSeq (sequencing and data preparation described in the methods)**.** We assembled this subset of reads with Geneious (default parameters) (Kearse *et al.* 2012) and used Blastn to find contigs with hits to mitochondrial genes (non-redundant database, e-value < 0.001). In our initial assembly we found a single, complete, assembled, circular contig ~16kb long with high confidence hits to all mitochondrial genes. In all following steps, we used this sequence as the fully assembled mitochondria. The mitochondrial genome was also included during polishing with Pilon for the seven iterations. We then used the MITOS online portal (Bernt *et al.* 2013) to annotate the 16191bp assembled and polished mitochondrial genome.

We attempted to assemble parts of the Y chromosome using sequencing information available for male *D. innubila* (SRA: SAMN07638923/SRR6033015) (Hill and Unckless 2017). We mapped these sequences to the female reference genome using BWA MEM with default parameters (Li and Durbin 2009), extracted all unmapped reads using SamTools (Li *et al.* 2009) and attempted to assemble these using Spades (default parameters) (Bankevich *et al.* 2012). We then mapped male and female expression data using GSNAP (Wu and Nacu 2010) to this dataset along with the whole genome. We considered the assembled contigs containing genes with significantly greater expression in males (using EdgeR (Robinson *et al.* 2009)), using the methods for RNA differential expression described below, $p$-value < 0.001, FDR < 0.001) to be putatively Y-linked. This filtered left us with 27 putatively male biased, Y-linked (or heterochromatic) scaffolds. We used blastn and tblastx to attempt to identify any known orthologs to these genes.

We identified large structural variants among the genomes of *D. innubila, D. falleni* and *D. phalerata* using both Manta (Chen *et al.* 2016) and Pindel (Ye *et al.* 2009) (default, bam input in both cases) on *D. falleni* and *D. phalerata* short read data mapped to the *D. innubila* genome. We extracted the structural variants found with both software packages as VCF files and considered only the variants detected by both Manta and Pindel to be real. We compared $dN/dS$ between genes found within inversions and outside and found no significant differences in either $dN/dS$ or $dS$ (Wilcoxon rank sum W = 35, $p$-value > 0.05).

For all genes we performed a codon bias analysis using CodonW (Peden 1997). We compared the codon bias index (CBI), codon adaptation index (CAI) and the frequency of optimal codons (Fop) across scaffolds, between novel genes and previously known genes, and between highly expressed genes (counts per million reads [CPM] > 1 in at least one dataset) and under expressed genes (CPM < 1 across all datasets). We find a significant conservation of codons in the mitochondria, Muller element F and the heterochromatic contigs, versus all other contigs (Supplementary Figure 4, Wilcox test W > 1132, $p$-value < 0.01186 for all CBI, CAI and Fop) (Zhou and Bachtrog 2015). For Muller elements A, B and E, we find significant levels of codon adaptation, optimal use and codon bias (Supplementary Figure 4, Wilcox test W > 14217000 $p$-value < 0.001586 for all CBI, CAI and Fop). We find significant positive associations between gene expression, and codon adaptation and optimization (GLM t-value > 4.746, $p$-value < 2.1e-06), consistent with an expectation for selection for codon efficiency in more highly expressed genes.

We find 393 orphan genes in the genome and compared the median expression, gene length, number of introns, codon bias and GC content between previously identified genes and the remaining putatively novel genes using codonW (Peden 1997) and bedTools (Quinlan and Hall 2010). Orphan genes are significantly shorter, under-expressed, AT-rich and intron-poor when compared to genes with previously identified orthologs (Supplementary Figure 5, Wilcoxon rank sum $p$-value < 0.0113), consistent with their more recent origin (Palmieri *et al.* 2014). We find a significant excess of orphan genes on two unassembled (scaffolds 5 and 11), these scaffolds are likely heterochromatic and sparse coding regions ($\chi^2 > 16.7$, $p$-value < 0.0005), We also find a significant deficit of orphan genes on Muller elements C and E ($\chi^2 >$ 14.17, $p < 0.000836$). Of these orphans, 51 show differential expression across life stages, primarily in the embryos, suggesting possible functionalization in different stages (Supplementary Tables 7-10, EdgeR analysis $p$-value < 0.05, FDR < 0.05 after multiple testing correction).

*Transposable element (TE) family comparison between species of the quinaria group*

We identified repetitive sequences *de novo* using RepeatModeler (engine = NCBI) (Smit and Hubley 2008). We then used RepeatMasker to mask the repetitive regions and classify repeats in classes/orders/families (-gff –gcalc –s) (Smit and Hubley 2015). We then used Blastn (parameters: $e$-value < 0.001) to compare each consensus sequence identified to the Repbase TE database (Bao *et al.* 2015), to confirm the TE order of each sequence . Using the GFF of repeat sequences generated by RepeatMasker, we then calculated the insertion density per 250kbp of the genome sliding across the genome for TE insertions. Using genomeCoverageBed (Quinlan and Hall 2010), we found the median coverage of the autosomes and each TE family and estimated the copy number of each TE family in the genome.

We estimate 13.53% of the genome consists of transposable elements (TEs). We find 175 TE families, consisting of 79 terminal inverted repeat DNA transposon families (TIR, 5.01%), 34 rolling circle/helitron DNA transposon families (RC, 5.61%), 25 long terminal repeat retrotransposons (LTR, 1.04%), and 26 long interspersed nuclear element retroposons (LINE, 1.87%) (Table 1). In addition to transposable elements, we find 10 short interspersed nuclear elements (SINE) and satellite element families, which together with simple repeats make up 3.42% of the genome (Figure 1A, Supplementary Figure 6). On Muller element A and B, we find two large regions consisting primarily of transposable elements. We considered these to be heterochromatic regions and potentially piRNA clusters. A majority (over 50% of the sequence) of these clusters consists of single TE superfamilies. Helitrons are primarily found throughout Muller element A, while Muller B's heterochromatic region consists of R2 LINE retroposons (Figure 1).

For *D. innubila, D. falleni* and *D. phalerata*, we mapped the short read information to the masked species reference genome with concatenated consensus TE sequences using BWA MEM (parameters: -t 4) (Li and Durbin 2009; Li *et al.* 2009). Following this we counted the proportion of reads mapping to each TE sequence of all reads, and the coverage of each TE sequence, weighted by the median coverage of the Muller element D. We removed all TE sequences with coverage for less than 80% of the sequence for less than 1x the median coverage of Muller element D, checked using bedTools GenomeCoverage (Quinlan and Hall 2010). In *D. innubila* we find 6136 TE copies, primarily TIR and RC DNA transposons (2688 and 2423 copies respectively). In *D. falleni* and *D. phalerata*, we see an expansion of LINE retroposons (1107 and 1793 copies respectively, versus 797 copies in *D. innubila*). We find a significant correlation between copy numbers of families for pairwise comparisons of all three species (Pearson's correlation = 0.51-0.68, $p$-value $< 2.42e-11$, $t$-value $> 7.174$), though specific families seem to differ wildly in copy numbers between species (Supplementary Figure 6C).

Finally we also used dnaPipeTE to get an independent estimate of the TE content (Goubert *et al.* 2015), using the *D. innubila* estimated genome size and the next generation sequencing information for each species (dnaPipeTE parameters: 2 iterations of trinity, 168Mb genome size, 1x estimated genome coverage reads). Comparing between species, we find even more dramatic differences, including a huge expansion of simple repeats in the *D. falleni* genome, accompanying an expansion of LINE elements, and an expansion of LTRs and TIRs in *D. phalerata* (Supplementary Figure 6C). Notably, these do not match the estimated TE proportions in Supplementary Figure 6B, it suggests *D. falleni* and *D. phalerata* contain TE families not present in *D. innubila*.

To identify TEs with orthology to known sequences, we used Blastn (parameters: -evalue 0.00001) against the Repbase Arthropod TE database (Bao *et al.* 2015). We grouped sequences with hits to previously identified TEs by the TE order and species family of the host. For 92 of the 175 TE families, we could identify a closely related TE sequence in a previously sequenced genome from RepBase (Supplementary Figure 7, Blastn, $e$-value $< 0.001$). Most these families (73.9%) are DNA transposons and LTRs, consistent with previous findings that these orders are more likely to be more recently horizontally transmitted, compared to LINEs (Bartolomé *et al.* 2009; Peccoud *et al.* 2017). 86 of these putatively horizontally transferred TE sequences are found in another *Drosophila* genome, with 6 TIR families with Blastn hits for Carpenter ants (*Camponotus*), likely found in the same environment as *D. innubila* (Patterson and Stone 1949; Markow and O'Grady 2006). Among the TEs with hits to *Drosophila*, only 32 (35.9%) are to *Drosophila* subgroup species thought to overlap in range with *D. innubila*, the remainder are species within the *Sophophora* subgroup (Supplementary Figure 7). While 54 of these TE families have hits to *Sophophora* species found in overlapping ranges with *D. innubila*, such as species in the *pseudoobscura*, *willistoni* and *ananassae* (within *melanogaster*) groups (Markow and O'Grady 2006), several TEs (20 TEs with hits to *melanogaster* group), show no evidence of this, with hits to species endemic to Asia or Africa (Supplementary Figure 7). This may be because these TEs share a common ancestor in the genome of an unsequenced species that has overlapping ranges with both *D. innubila* and the *melanogaster* group species.

*Identifying duplications*

We identified the 1014 genes present in multiple copies in *D. innubila*, but only present as single copies in *D. virilis* and *D. melanogaster*. Most these (866) have the duplicated copy on the same chromosome, with most these duplicates (848) within 50kb of the original copy (determined by the position of the ortholog in *D. melanogaster*). These duplications are enriched for metal ion transport and protein metabolism genes (GOrilla, $p$-value < 0.0005, FDR < 0.05, enrichment > 1.65) (Eden *et al.* 2009), including 26 cytochrome P450 recent duplications. For each set of duplicates, we extracted the coding sequence and aligned using PRANK (-codon +F –f=paml) (Löytynoja 2014). We identified positive selection between orthologs using codeML, for models M0, M1a, M2a and M3 (Yang 2007). We used a likelihood ratio test to identify which model fits best for each set of orthologs and to identify duplicates under putative adaptive evolution. Of duplicate genes, 294 (28.9%) showed signatures of positive selection, a higher proportion than seen in non-duplicated genes (4.7% , Supplementary Figure 8, $dN/dS > 1$, Model 2a is best fitting model). We find no association between the number of copies of a gene and the $dN/dS$ (GLM, $t$-value = 0.27, $p$-value = 0.78) and as shown previously, find negative correlations between $dN/dS$ and both gene length and dS (Supplementary Table 11, Supplementary Figure 8, GLM, $t$-value < -2.2353, $p$-value < 0.0188). Using GOrilla we found that, like the total complement of paralogs, these duplicated genes under positive selection are again enriched for Metal ion transport, specifically the copper ion response pathways (Supplementary Figure 8, Supplementary Table 11, GOrilla, *p*-value < 0.0005, FDR < 0.05, enrichment > 1.25) (Eden *et al.* 2009).

*RNA differential expression analysis*

We downloaded mapped RNA sequencing information from ModEncode (modencode.org) for *D. melanogaster* across all life stages (Chen *et al.* 2014).

For each set of *D. innubila* RNA sequencing short read information we mapped it to the masked *D. innubila* genome with the TE sequences concatenated to the end using GSNAP. We then counted the number of reads mapped to each gene per kb of gene using HTSeq for all mapped RNA sequencing data and normalized by counts per million per dataset (Anders *et al.* 2015).

We then used the R package EdgeR (Robinson *et al.* 2009) to make differential expression comparisons between the following datasets: 1. Adult total body *D. innubila* RNA, male versus female; 2. RNA across different life stages total body; 3. Adult total body, *D. innubila* female versus *D. melanogaster* female; 4. Adult total body, *D. innubila* male versus *D. melanogaster* male; 5. Adult total body, *D. innubila* versus *D. melanogaster*; 6. Larvae total body, *D. innubila* versus *D. melanogaster*;  7. Pupae total body, *D. innubila* versus *D. melanogaster*; 8. Whole embryo, *D. innubila* versus *D. melanogaster*. In each case, we compared the counts per million per 1kbp exon of genes to identify significant differences in expression of orthologous genes (*p*-value < 0.05, FDR < 0.05 after adjusting for multiple testing).

Following this, we used GOrilla (Eden *et al.* 2009) to identify and visualize enriched gene ontology (GO) terms, separating by genes that are and aren't differentially expressed (p-value threshold = 0.001) for process, function and component GO terms. For functional terms of interest, such as detoxification genes between species, recent duplications versus their single copy, novel genes across life stages or viral RNAi genes, we compared expression differences between groups by hand. Across the life stages between *D. innubila* and *D. melanogaster*, we find changes in gene expression, including enrichments such as muscle system process genes and structural muscle construction. We also find differential expression metabolic processes, cellular process and locomotion across all life stages (Supplementary Tables 12-19, Supplementary

Figure 9, GOrilla, FDR < 0.00005, $p$-value < 0.000984 after multiple testing correction, enrichment > 1.21), it is important to highlight that these differences identified could be due to differences in experimental setting used to generate the data, or could be due to differences between *D. innubila* and *D. melanogaster*.

Using our gene expression data for both male and female adult *D. innubila*, we looked for biases expected between sexes. Surprisingly, we find no genes with a significant female bias expression (0 genes, Supplementary Figure 10, Supplementary Table 7, 13 & 20, EdgeR $p$-value >0.206 FDR > 0.0006 after Bonferroni multiple testing correction), with a large number showing a male bias (Supplementary Figure 10, 223 genes, EdgeR p < 0.000001 after multiple testing correction). As is expected there is a significant deficit of male bias genes on the X chromosome (Supplementary Tables 7 & 20, Chi-Square test $\chi^2$=4.21, $p$-value = 0.04), though we also see an enrichment on one of the autosomes, Muller element B (Chi-Square test $\chi^2$ = 16.86, $p$-value = 4.03e-5). We used GOrilla to identify any enrichment in categories between sexes which may explain the difference observed. We find an enrichment for organophosphate metabolism, cell motility and sperm movement (GOrilla enrichment > 17.27, $p$-value < 0.000654).

*Structural variants between species in the D. innubila trio*

We next estimated structural variants between *D. innubila*, *D. falleni* and *D. phalerata,* using Pindel (Ye *et al.* 2009) and short reads mapped to the *D. innubila* genome. We find many more structural variants and inversions between *D. phalerata* and *D. innubila* than *D. falleni,* consistent with structural variants accumulating as species diverge (Supplementary Figures 11 & 12). We find no significant effects of inversions on *dN/dS* or *dS* between species (Mann-Whitney U test W < 156, $p$-value > 0.41).

**Supplementary Tables and Figures**

**Supplementary Table 1:** Summary of reads used for genome sequencing, assembly, annotation and *dN*/*dS* calculation.

**Supplementary Table 2:** Summary statistics for each iteration of the genome.

**Supplementary Table 3:** Summary of the genic characteristics of the *D. innubila* genome.

**Supplementary Table 4:** Genes ontologies (GO) enriched for genes with high/low residuals for *dN/dS* between *D. melanogaster* and *D. innubila*, due to drastic differences between the species.

Enriched categories are categories which are slow evolving in one species, but fast evolving in the other.

**Supplementary Table 5:** Summary of *dN*/*dS* statistics for each immune gene category across the total group and on each branch. Additionally, the t-score and p-value for a two-sided *t*-test ($\mu = 0$) for that category is shown. Significant categories are highlighted in bold.

**Supplementary Table 6:** *dN*/*dS* enrichment for *Drosophila innubila* trio for processes, components and functions, including any enrichments for specific branches.

**Supplementary Table 7:** GO enrichment for processes, components and functions for differential expression between *D. innubila* males and females.

**Supplementary Table 8:** GO enrichment for processes, components and functions for differential expression between *D. innubila* embryos and larvae.

**Supplementary Table 9:** GO enrichment for processes, components and functions for differential expression between *D. innubila* larvae and pupae.

**Supplementary Table 10:** GO enrichment for processes, components and functions for differential expression between *D. innubila* pupae and adults.

**Supplementary Table 11:** *dN*/*dS* GO enrichment for duplications for processes, components and functions, including any enrichments for specific branches.

**Supplementary Table 12:** A table summarizing the differential gene expression shown in Supplementary Tables 13-19, showing the number of genes differentially expressed between *D. innubila* and *D. melanogaster* at differing life stages, with enrichments in gene ontology (GO) categories.

**Supplementary Table 13:** GO enrichment for processes, components and functions for differential expression between *D. melanogaster* and *D. innubila* embryos.

**Supplementary Table 14:** GO enrichment for processes, components and functions for differential expression between *D. melanogaster* and *D. innubila* larvae.

**Supplementary Table 15:** GO enrichment for processes, components and functions for differential expression between *D. melanogaster* and *D. innubila* pupae.

**Supplementary Table 16:** GO enrichment for processes, components and functions for differential expression between *D. melanogaster* and *D. innubila* adults.

**Supplementary Table 17:** GO enrichment for processes, components and functions for differential expression between *D. melanogaster* and *D. innubila* adult males.

**Supplementary Table 18:** GO enrichment for processes, components and functions for differential expression between *D. melanogaster* and *D. innubila* adult females.

**Supplementary Table 19:** GO enrichment for processes, components and functions for differential expression between *D. melanogaster* and *D. innubila* total samples.

**Supplementary Table 20:** Enrichment or depletion of genes differentially expressed between male and female samples on each scaffold/Muller element including the $\chi^2$ for this enrichment.

**Supplementary Figure 1:** Histograms of dN/dS for *D. innubila* and *D. melanogaster*.

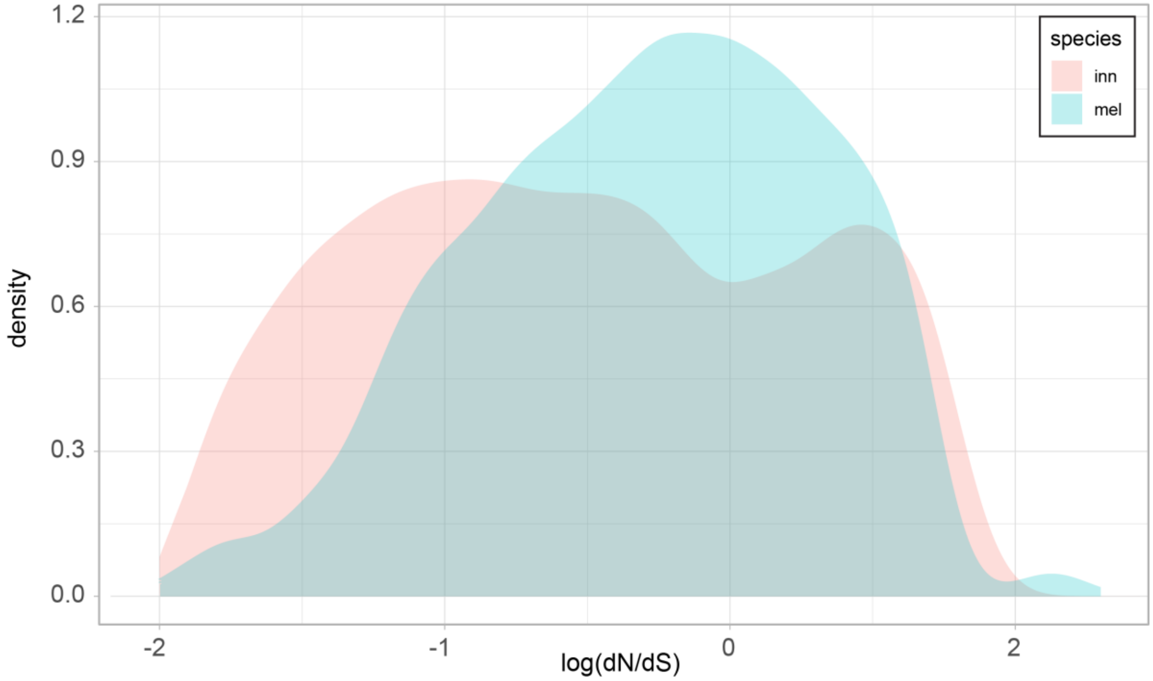


**Supplementary Figure 2:** δ (calculated using HyPhy) by immunity category for both *D. innubila* and *D. melanogaster*.

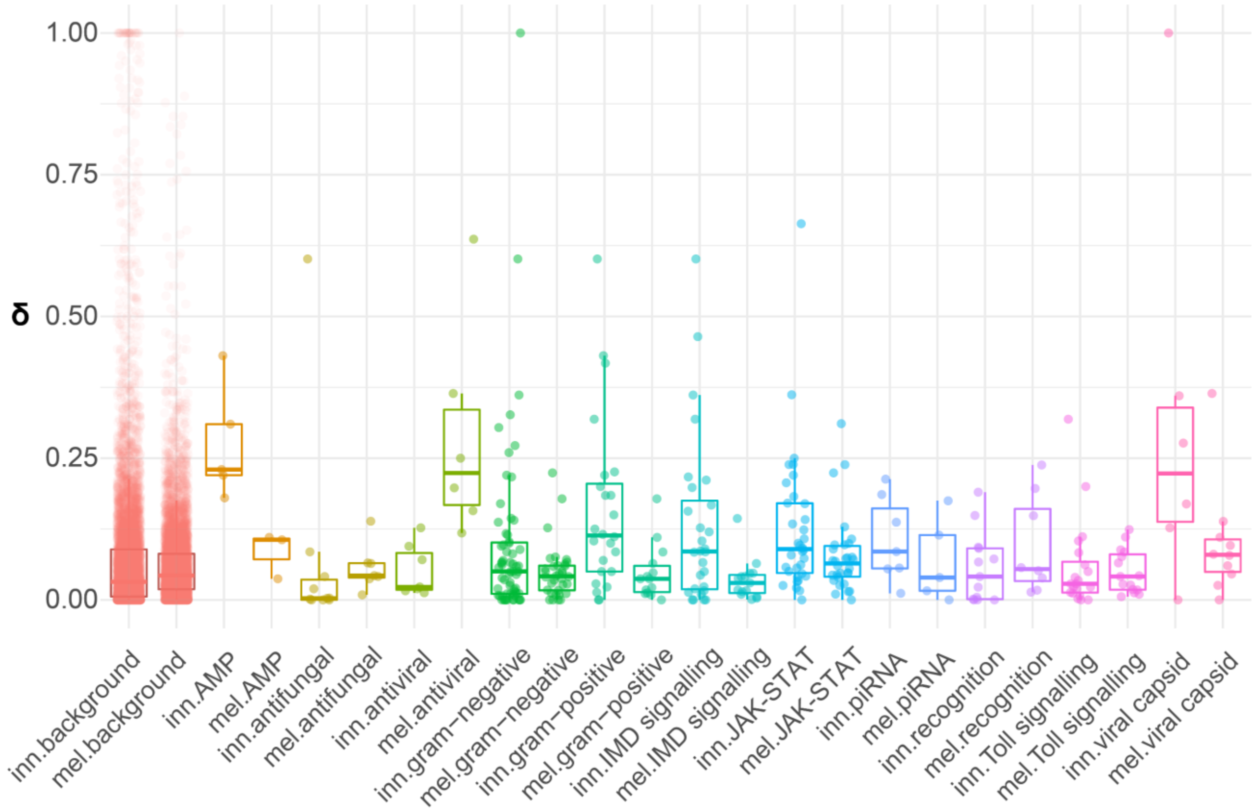

**Supplementary Figure 3: Antiviral evolution across the *quinaria* group.** Difference between viral RNAi, JAK-STAT (filled dots = regulatory, empty dots = cytokines), NF-κB, Toll and putatively viral-interacting proteins from the background *dN*/*dS* of genes of similar *dS* (+-0.01*dS*) for the *D. melanogaster* branch, the *D. innubila* branch, the total *D. melanogaster* tree and the total *D. innubila* trio. Genes known to be associated with the immune response to viral infection, but no known pathway are classed as 'Other Antiviral'. A *p*-value (from a two-sided *t*-test looking for significant differences from 0) of 0.05 or lower is designated with *.

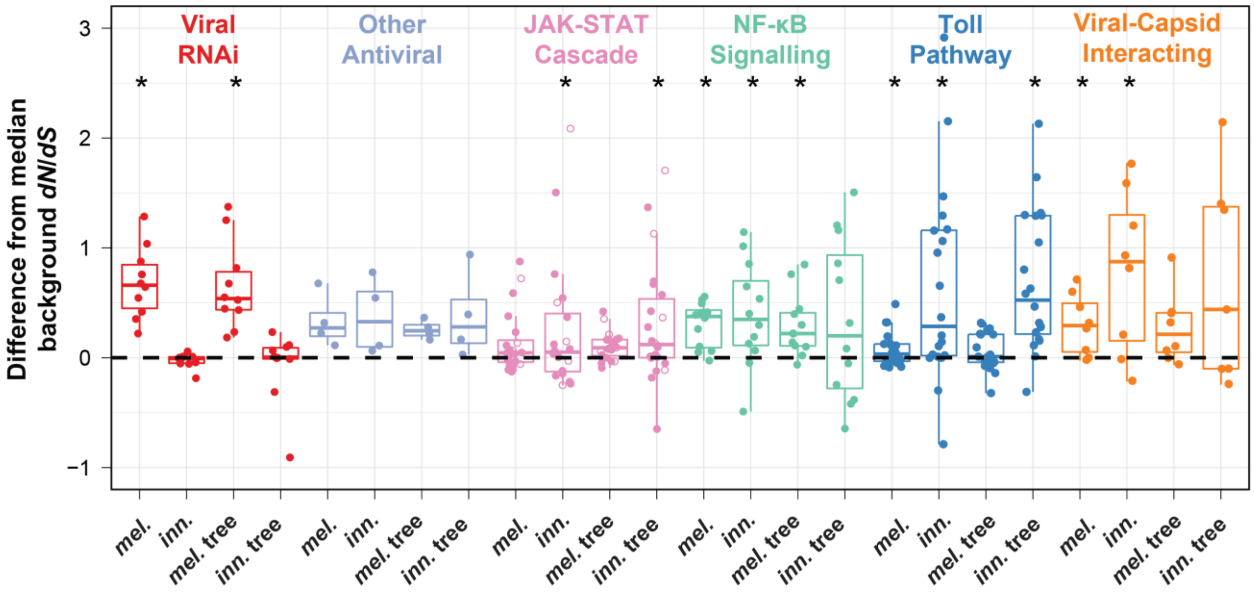

**Supplementary Figure 4:** Codon bias distributions across the *Drosophila innubil*a genome, separated by scaffold. CAI = Codon adaptation index. CBI = Codon bias index. Fop = Frequency of optimal codons. GC = Proportion of GC across each gene.

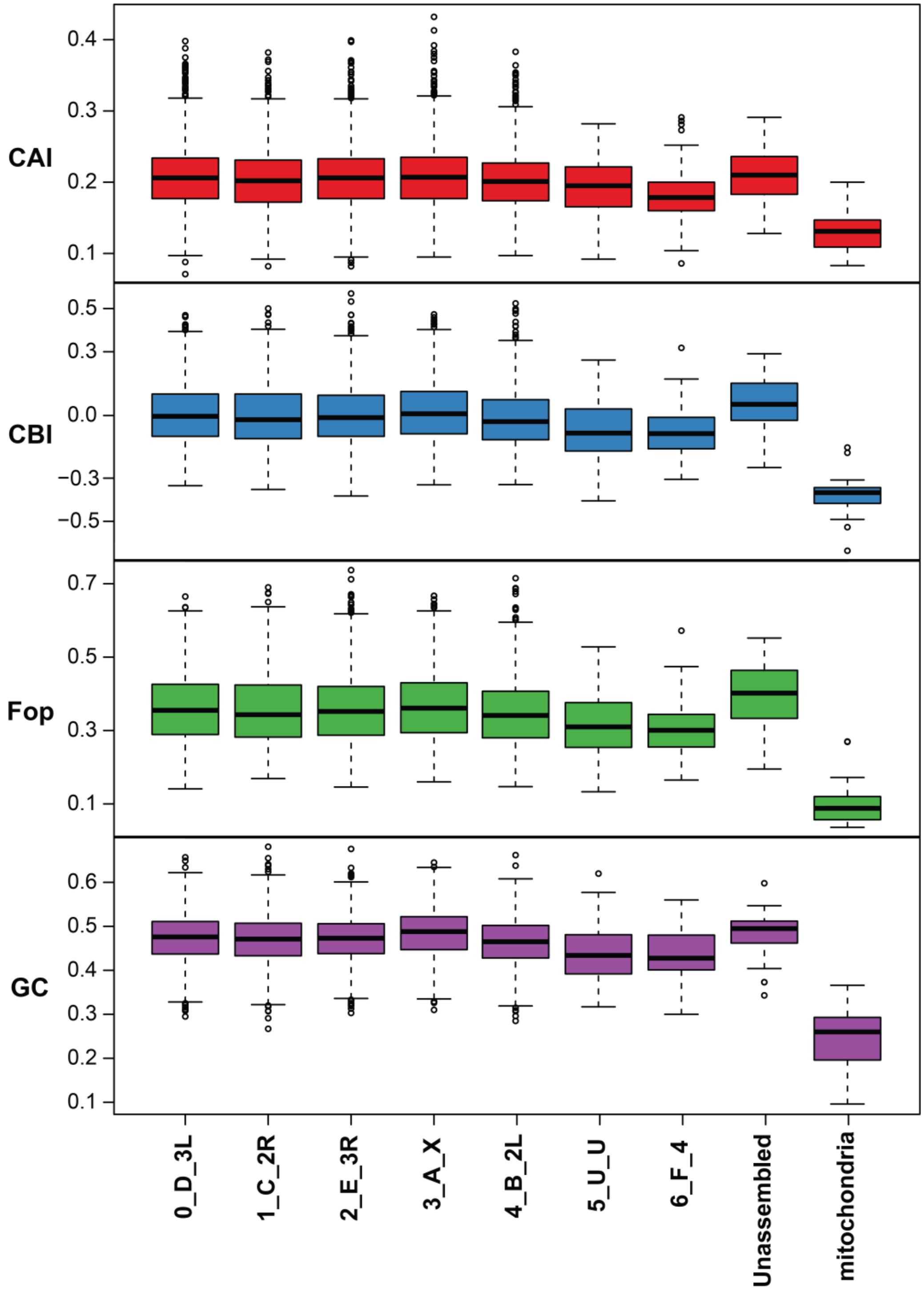

**Supplementary Figure 5:** Comparison between orphan genes and previously described genes, including: **A.** Codon adaptation index (CAI). **B.** Codon bias index (CBI). **C.** Frequency of optimal codons (Fop). **D.** Gene length (in bp). **E.** Number of introns per gene. **F.** Mean expression across life stages (read counts per million).

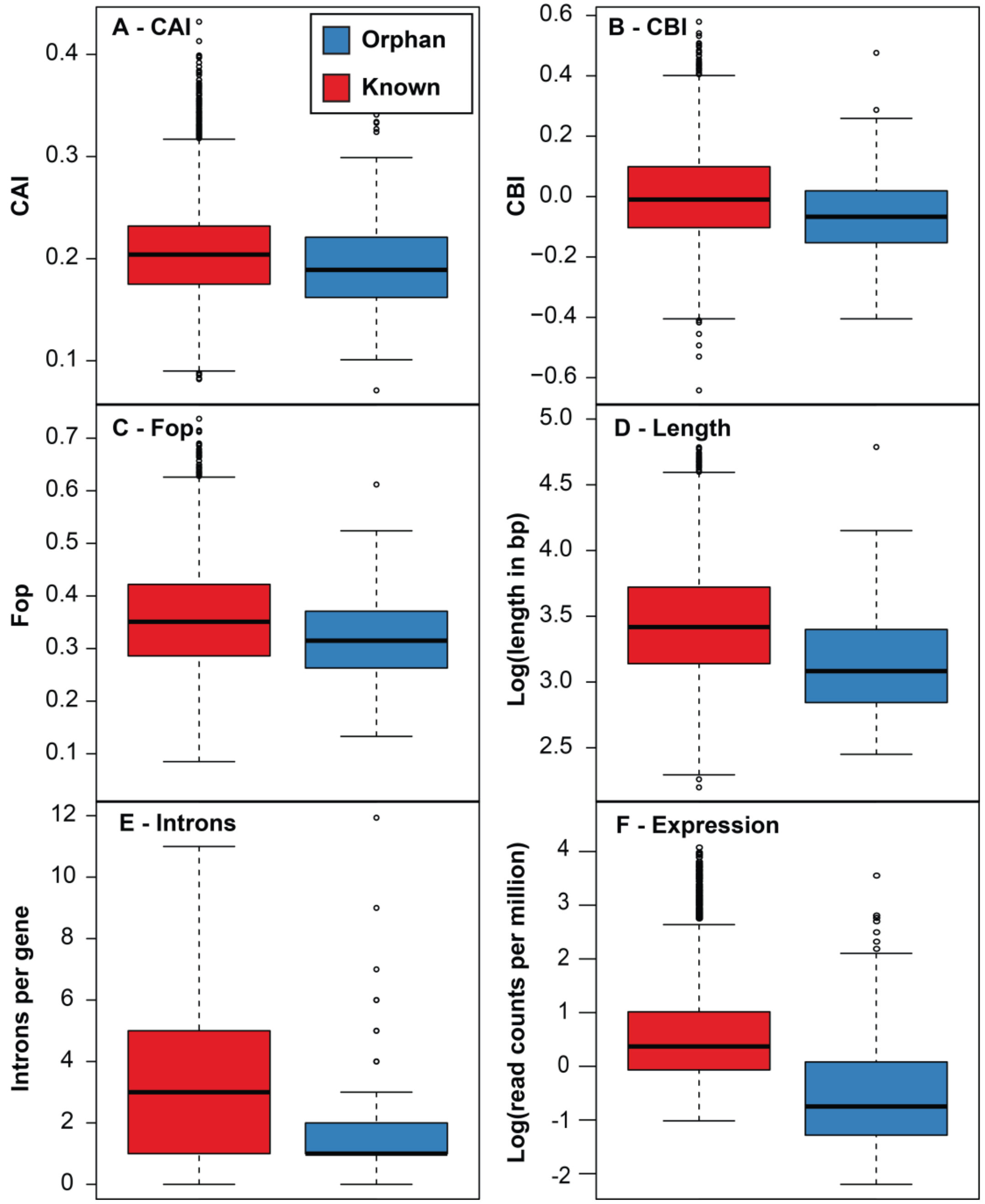

**Supplementary Figure 6: A.** The proportion of the *D. innubila* genome masked by each type of repeat. LINE = Long interspersed nuclear element RNA transposon, LTR = long terminal repeat RNA transposon, RC = rolling circle DNA transposon, TIR = terminal inverted repeat DNA transposon. **B.** TE content of *D. innubila*, *falleni* and *phalerata,* **C.** Copy number comparisons between *D. innubila*, *D. falleni* and *phalerata*. **D.** dnaPipeTE estimates of the genomic proportion of repetitive elements for each species examined. Other, NA and SINE categories were removed due to small proportions. Though unlabeled, rRNA is shown in yellow and constitutes 1-2% of the genome.

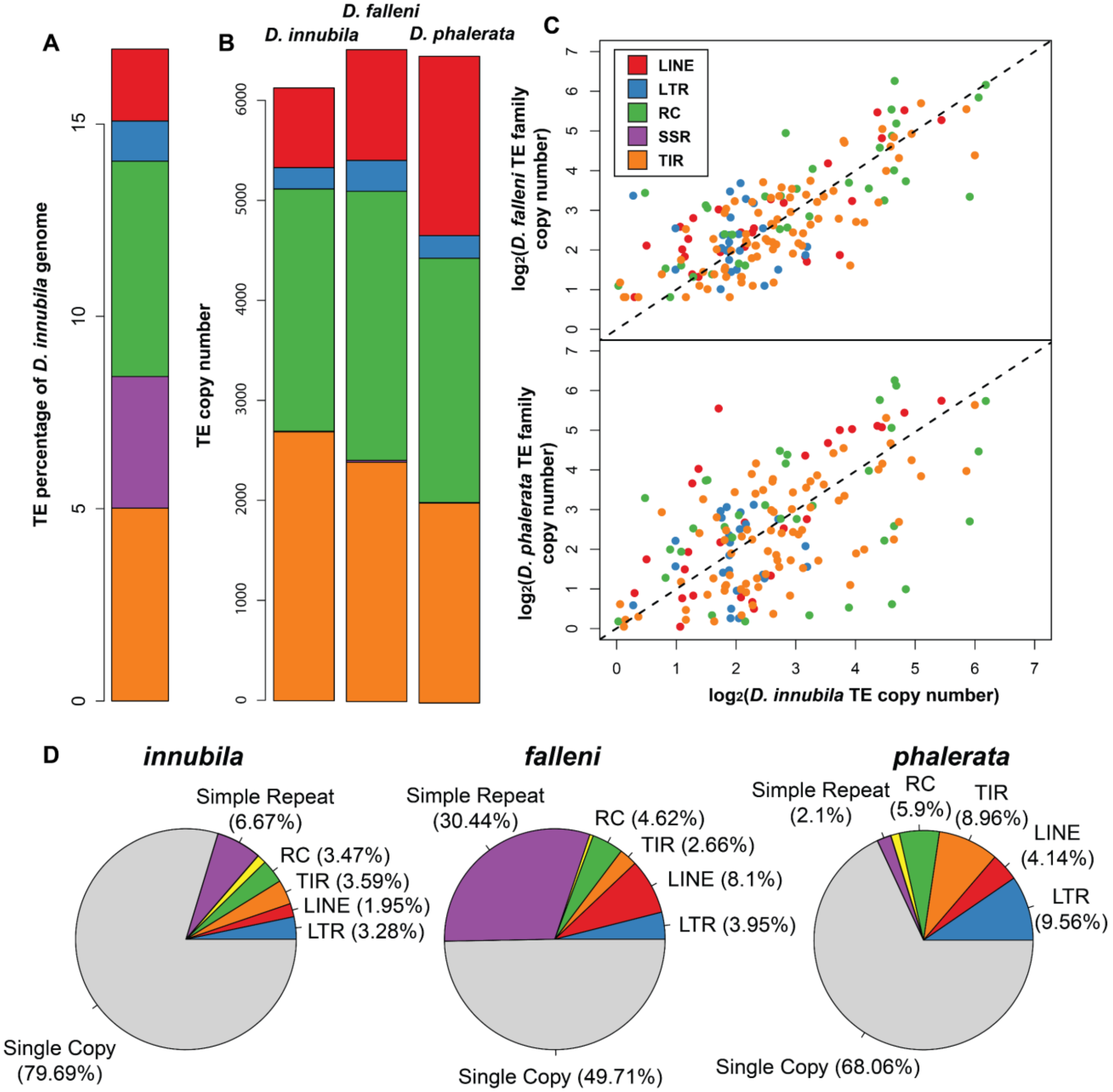

**Supplementary Figure 7:** Number of TE families found in *D. innubila,* closely related to known TE families (taken from Repbase) in different species group, identified using BLAST, suggesting relatively recent horizontal transfer events.

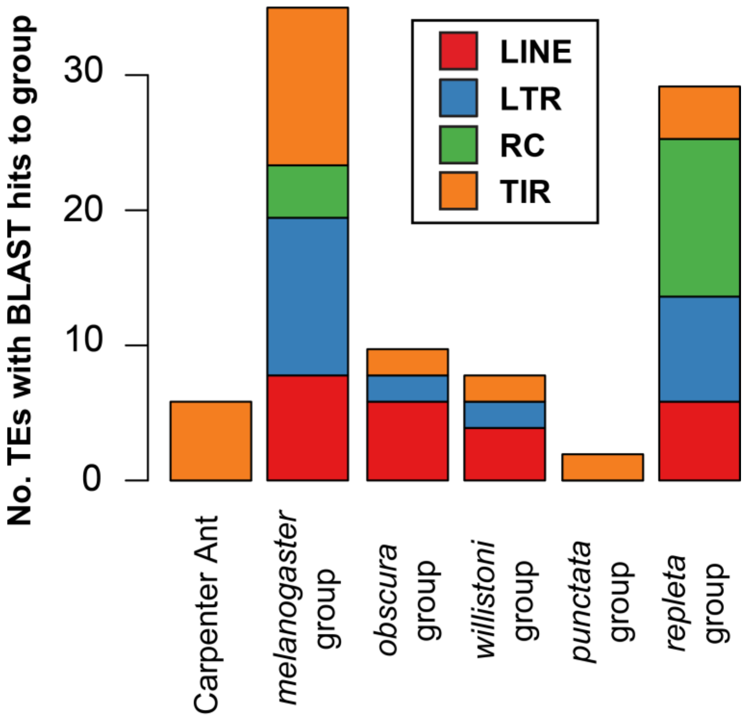

**Supplementary Figure 8:** *dN/dS* versus *dS* across paralogs for recently duplicated genes. Metal ion binding, protein metabolism and immunity genes are highlighted.

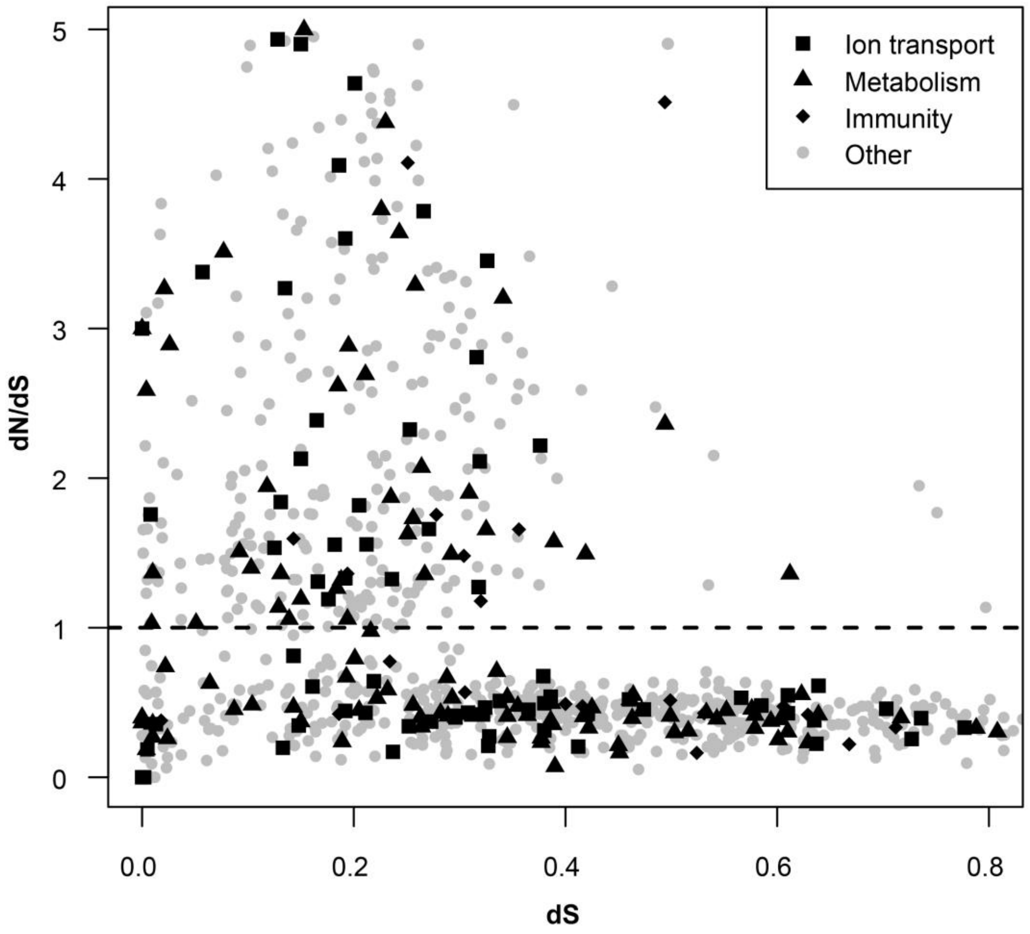

**Supplementary Figure 9:** Volcano plots sowing differential gene expression between *D. innubila* and *D. melanogaster* at different life stages. Dots are colored by their significance and if a recent duplication or not (duplicants layered on top), the significance cut off is set a 0.05 following Bonferroni multiple testing correction.

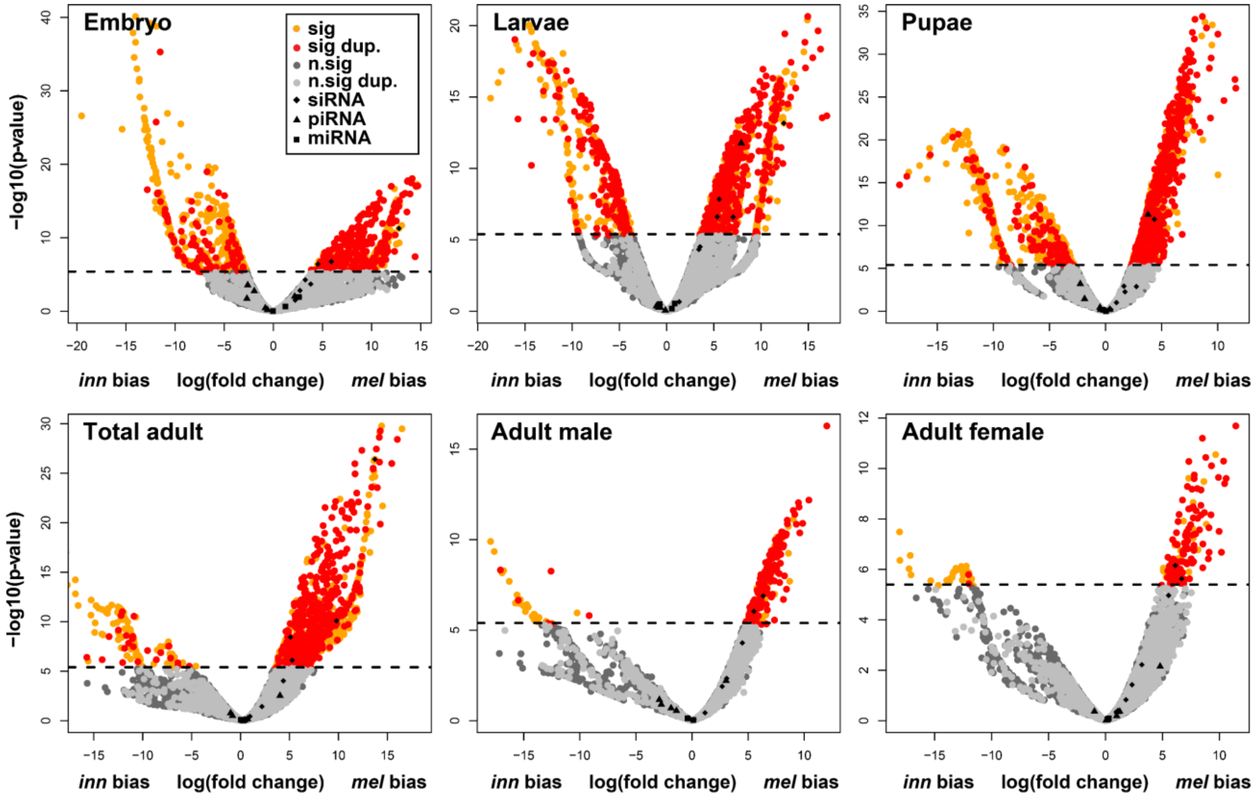

**Supplementary Figure 10:** Volcano plot showing differential gene expression between *D. innubila* male and female samples and significant differences, highlighting if genes are duplicated relative to *D. virilis* or not, and if genes are involved in sperm motility.

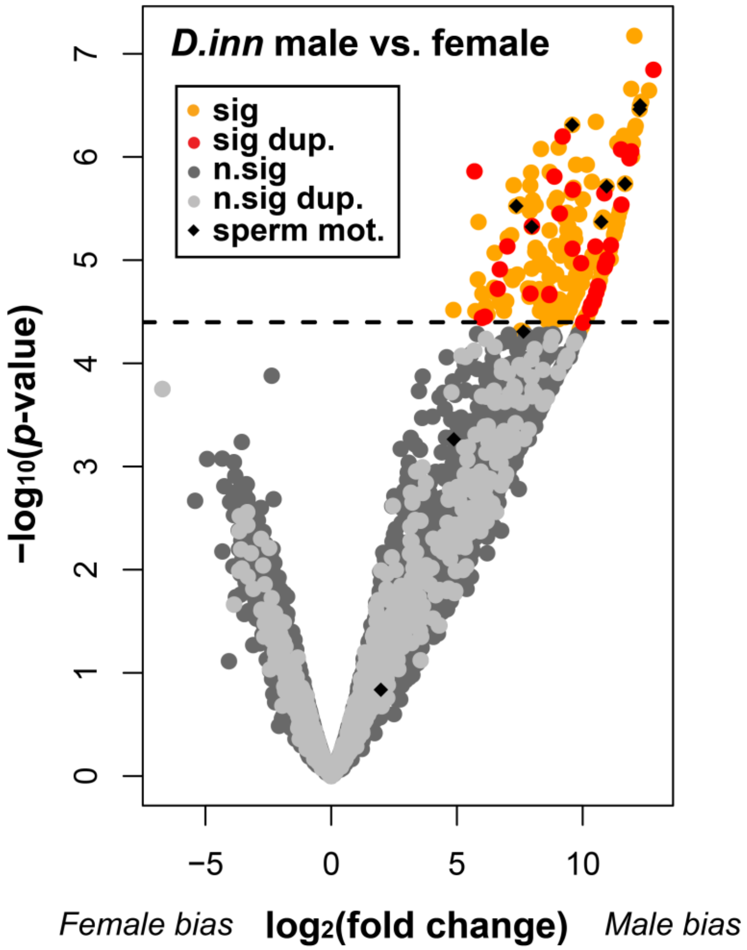

**Supplementary Figure 11:** Inversions identified between *D. innubila and D. falleni, and between D. innubila/falleni and D. phalerata* using Pindel (Ye et al. 2009) and Manta (Chen et al. 2016) (taking the consensus of the two programs). Scaffolds are labelled and colored by the Muller element they belong to.

**A - *falleni* inversions** **B - *phalerata* inversions**

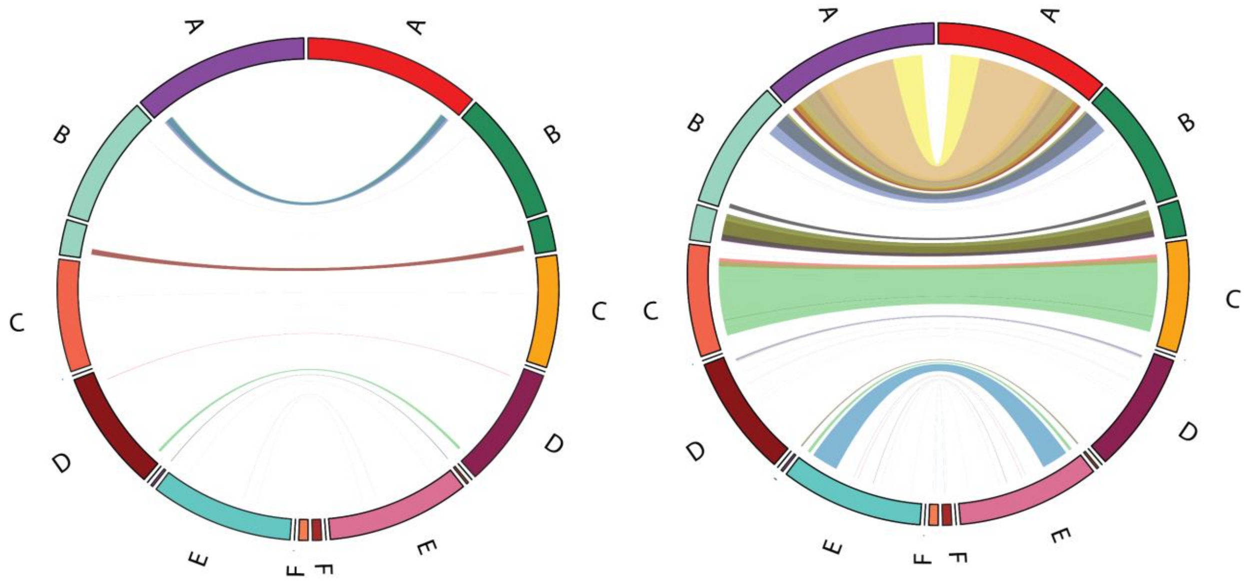

**Supplementary Figure 12:** Size and number of each structural variant between *D. innubila* and *D. falleni* identified using Pindel and Manta (taking the consensus of the two programs).

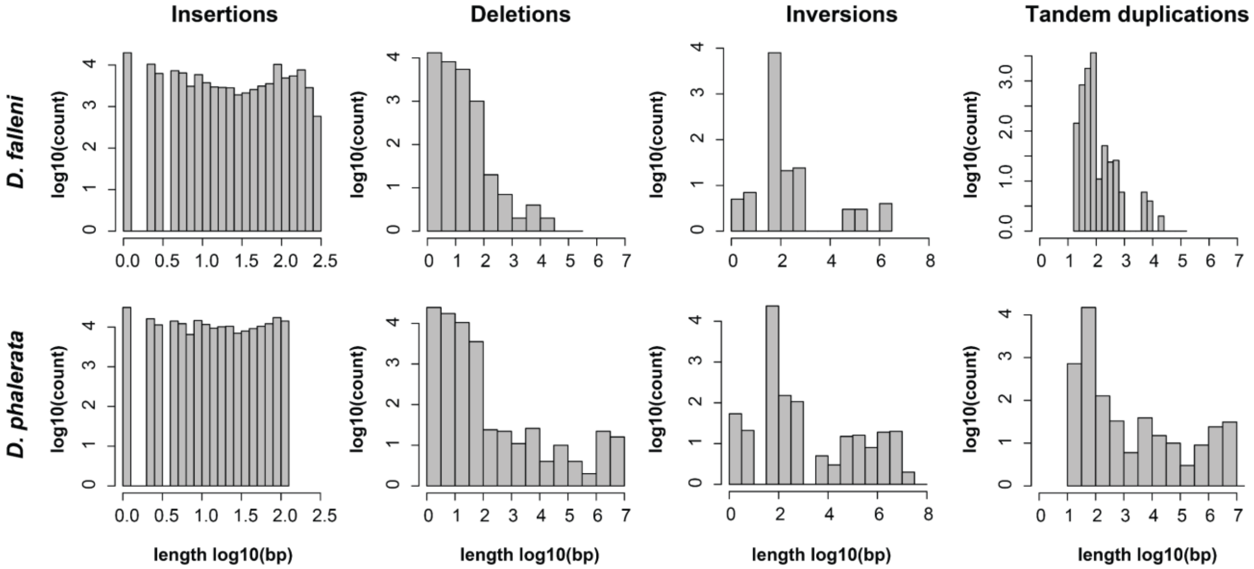
